# MEIG1/PACRG associated and non-associated functions of axonemal dynein light intermediate polypeptide 1 (DNALI1) in mammalian spermatogenesis

**DOI:** 10.1101/2022.04.28.489920

**Authors:** Yi Tian Yap, Wei Li, Qian Huang, Qi Zhou, David Zhang, Ljljiana Mladenovic-Lucas, James G Granneman, David C Williams, Rex A Hess, Aminata Touré, Zhibing Zhang

**Affiliations:** Department of Physiology, Wayne State University School of Medicine, Detroit, MI, 48201; Department of Occupational and Environmental Medicine, School of Public Health, Wuhan University of Science and Technology, Wuhan, Hubei, 430060, China; College of William and Mary, Williamsburg, VA, 23187, USA; Center for Molecular Medicine and Genetics, Wayne State University School of Medicine, Detroit, MI, 48201, USA; Department of Pathology and Laboratory Medicine, University of North Carolina, Chapel Hill, NC 27599; Department of Comparative Biosciences, College of Veterinary Medicine, University of Illinois, 2001S. Lincoln, Urbana, IL 61802-6199; Institut pour l’avancée des Biosciences, Université Grenoble Alpes, INSERM U1209, CNRS UMR 5309, F-38000 GRENOBLE, France; The C.S. Mott Center for Human Growth and Development, Department of Obstetrics & Gynecology, Wayne State University, Detroit, MI, 48201

**Keywords:** DNALI1, manchette, cargo transport, spermiogenesis

## Abstract

Axonemal dynein light intermediate polypeptide 1 (DNALI1) was originally cloned from *Chlamydomonas reinhardtii* in an effort to find motor proteins essential for flagellar motility. Here we report that DNALI1 is a binding partner of parkin co-regulated gene 1 (PACRG), which forms a complex with meiosis expressed gene 1 (MEIG1) in the manchette, a transient and unique structure only present in the elongating spermatids and required for normal spermiogenesis of the male germ cell differentiation process. DNALI1 recruits the PACRG protein in transfected CHO cells, and also stabilizes PACRG in bacteria and transfected mammalian cells. The untagged DNALI1 could also be co-purified with His-tagged PACRG in the gel filtration assay. Immunofluorescence staining on isolated male germ cells revealed that DNALI1 was present in the manchette of elongating spermatids, and colocalized with PACRG in this structure. In *Pacrg* mutant mice, localization of DNALI1 in the manchette was not changed, suggesting that DNALI1 and PACRG form a complex in the manchette, with DNALI1 being an upstream molecule. Mice deficiency in DNALI1 specifically in male germ cells showed dramatically reduced sperm numbers and were infertile. In addition, majority of the sperm exhibited abnormal morphology including misshapen heads, bent tails and enlarged midpiece, discontinuous accessory structure, and loss of sperm individualization, emphasizing the importance of DNALI1 in sperm development. Examination of testis histology revealed impaired spermiogenesis in the conditional *Dnali1* knockout mice. Electron microscopy revealed disrupted ultrastructure in sperm of the *Dnali1* mutant mice. Testicular levels of MEIG1, PACRG and SPAG16L proteins were not changed in the *Dnali1* mutant mice. However, MEIG1 and SPAG16L were no longer present in the manchette in the absence of DNALI1. These findings demonstrate that DNALI1 is involved in the connection of the MEIG1/PACRG complex to carry cargo proteins along the manchette microtubules for sperm flagella formation. Given that *Dnali1* mutant mice showed impaired sperm individualization that was not observed in the MEIG1 nor PACRG-deficient mice, DNALI1 might fulfill other functions beyond its role associated with the MEIG1/PACRG complex. Thus, DNALI1 plays multiple roles in sperm cell differentiation and function.

**Summary statement:** Axonemal dynein light intermediate polypeptide 1 (DNALI1) is required for sperm formation and male fertility. It associates with the MEIG1/PACRG complex in the manchette and is involved in a cargo transport system. In addition, it might be related to IFT and sperm individualization.

## Introduction

Motile cilia are microtubule-based organelles with a core “9 + 2” axonemal structure (1). They generate fluid flow by their beating in various epithelia and also enable sperm cell progression, which overall confers vital functions in eukaryotes (2, 3). In humans, motile cilia dysfunction results in multiple syndromes, including bronchiectasis, impaired mucociliary clearance, chronic cough, sinusitis, and male infertility, which are described as immotile cilia syndrome, as well as *primary ciliary dyskinesia* (PCD) (4, 5). Ciliary beat is driven by dynein motor protein complexes, associated with the inner and outer dynein arms of axonemal ultrastructure, which are anchored to the peripheral axonemal microtubules. The heavy, medium and light chains constitute the dynein motors, and each chain has a different molecular weight and provides an important function (6, 7). Hence, in mammals the loss of dynein function leads to PCD with ciliary impairment with hydrocephalus, body axis asymmetry, and male infertility (2, 8, 9).

Human axonemal dynein light intermediate polypeptide 1 (*DNALI1*) is the homolog of the *Chlamydomonas* inner dynein arm gene *p28*, an important component of the ciliated dynamic arm, which is mainly responsible for cilium movement (10, 11). The molecular analysis of *DNALI1 (*human *hp28*) gene was revealed by Dr. Shalender Bhasin’s laboratory more than 20 years ago. The gene was localized on chromosome 1 region p35.1, and *DNALI1* transcripts were detected in several ciliated structures, including sperm flagella, suggesting that this gene could be a good candidate for patients suffering from immotile cilia syndrome (11). In addition to PCD, *DNALI1* was also related to a variety of diseases, including Frontotemporal Lobar Degeneration (FTLD), Diploid Breast Carcinoma (DPC), Osteosarcoma (OS), Allergic Rhinitis (AR), Nasopharyngeal Carcinoma (NPC), and Alzheimer’s disease (AD) (12–18), although the precise mechanisms are unclear. However, the role of *DNALI1* in human male reproduction has not been investigated.

Recently, *Dnali1* was identified as a sex-related gene involved in spermatogenesis of several fishes, including the Odontobutis potamophila, the Ussuri catfish Pseudobagrus ussuriensis, and the olive flounder Paralichthys olivaceus (19–21). The expression of *Dnali1* showed sexually dimorphism with predominant expression in the testis. This suggested that the fish *Dnali1* might play an important role in the testis, especially in the period of spermatogenesis (21). Sajid et al. cloned the murine *Dnali1* with significant similarity to the *p28* gene of *Chlamydomonas reinharditii* and to *DNALI1* (*hp28*) gene (22). The murine *Dnali1* gene is localized on chromosome 4 and consists of six exons. It has two transcripts and is expressed in several tissues, but the strongest expression was observed in testis. During the first wave of spermatogenesis, both *Dnali1* mRNA and protein were dramatically increased during the spermiogenesis phase, which corresponds to spermatid cell differentiation. Immunofluorescence studies demonstrated that DNALI1 was detected in spermatocytes and abundant in round and elongated spermatids. Moreover, the DNALI1 protein was localized in flagella of mature sperm (22), indicating that *Dnali1* may play an essential role in mouse sperm formation and male fertility.

Sperm production and formation is a complex process consisting of mitosis, meiosis and finally spermiogenesis (23). During spermiogenesis, germ cells undergo dramatic changes with the formation of sperm-unique structures, including the flagellum (24). Many genes have been described to precisely regulate these morphologic changes (25). We previously discovered that mouse meiosis-expressed gene 1 (MEIG1) and Parkin co-regulated gene (PACRG) form a protein complex in the manchette, a transient microtubule structure localized in the head compartment of sperm cells during differentiation. It was demonstrated that this protein complex controls sperm flagellum formation, with PACRG being an upstream molecule of MEIG1 (26–29). The MEIG1/PACRG complex was shown to function as part of the cargo transport system, called intra-manchette transport system (IMT) (30), which transports sperm flagella proteins, including SPAG16, for sperm tail assembly. However, how the MEIG1/PACRG complex associates with the axonemal motor system for cargo transport remained unclear.

Using mouse PACRG as bait for a yeast two-hybrid screen, DNALI1 was identified as a protein binding-partner. To investigate the function of DNALI1, a *Dnali1* conditional knock-out mouse model was generated using a floxed *Dnali1* mouse line, which was crossed with *Stra8*-iCre transgenic mice, to specifically disrupt the *Dnali1* gene in the male germ cells. Reported here, the *Dnali1* conditional knockout (*Dnali1* cKO) mice have significantly reduced sperm number and are infertile. In addition, the majority of sperm exhibited abnormal morphology, strongly indicating the importance of *Dnali1* in the development of sperm flagella.Importantly, in the conditional *Dnali1* mutant spermatid germ cells, MEIG1 and SPAG16L were not present in the manchette, indicating that DNALI1 is required for MEIG1/PACRG complex localization and for transport of cargo proteins to assemble the sperm flagellum. Some unique phenotypes discovered in the *Dnali1* mutant mice but not in the MEIG1 and PACRG-deficient mice suggest additional roles of the gene in sperm cell differentiation.

## Materials and methods

### Ethics statement

All animal research was executed in compliance with the guidelines of the Wayne State University Institutional Animal Care with the Program Advisory Committee (Protocol number: 18-02-0534).

### Yeast two-hybrid experiments

Full-length mouse PACRG coding sequence was amplified using the following primers: forward: 5’-<u>GAATTC</u>ATGCCGAAGAGGACTAAACTG-3’; reverse: 5’-<u>GGATCC</u>GTCAGTTCAGCAAGCACGACTC-3’. After TA cloning and sequencing, the correct cDNA was subcloned into the EcoR1/BamH1 sites of pGBKT7, which was used to screen a Mate & Plate™ Library-Universal Mouse (Normalized) (Clontech, Mountainview, CA; Cat#: 630482) using the stringent protocol according to the manufacturer’s instructions. For direct yeast two-hybrid assay, the coding sequence of the DNALI1 cDNA was amplified by RT-PCR using the following primers: forward: 5’-<u>GAATTC</u>ATGATACCCCCAGCAGACTCTCTG-3’ and reverse: 5’-<u>GGATCC</u>GATCACTTCTTCGGTGCGATAATGCC-3’. The correct cDNA was cloned into EcoRI/BamH1 sites of pGAD-T7 vector. The yeast was transformed with the indicated plasmids using the Matchmaker™ Yeast Transformation System 2 (Clontech, Cat#: 630439). Two plasmids containing simian virus (SV) 40 large T antigen (LgT) in pGADT7 and p53 in pGBKT7 were co-transformed into AH109 as a positive control. The AH109 transformants were streaked out in complete drop-out medium (SCM) lacking tryptophan, leucine and histidine to test for histidine prototrophy.

### Localization assay

To generate mouse PACRG/pEGFP-N_2_ plasmid, *Pacrg* cDNA was amplified using the primer set: forward: 5’-<u>GAATTC</u>ATGCCGAAGAGGACTAAACTG-3; reverse: 5’-<u>GGATCC</u>GGTTCAGCAAGCACGACTC-3’, and the correct *Pacrg* cDNA was ligated into the pEGFP-N_2_ vector. To generate mouse DNALI1/Flag construct, *Dnali1* cDNA was amplified using the primer set: forward: 5’-<u>GAATTC</u>AATGATACCCCCAGCAGACTCTCTG-3’; reverse: 5’-<u>CTCGAG</u>TCACTTCTTCGGTGCGATAATGCC-3’, and the correct *Dnali1* cDNA was ligated into the pCS3+FLT vector. The PACRG/pEGFP-N_2_ and DNALI1/Flag were transfected individually or together into CHO cells by using Lipofectamine™ 2000 transfection reagent (Invitrogen, Waltham, MA). The CHO cells were cultured with DMEM (with 10% fetal bovine serum) at 37°C. After 48 h transfection, the CHO cells were processed for immunofluorescence with an anti-Flag antibody.

### Co-immunoprecipitation assay

Mouse *Pacrg* cDNA was amplified using the following primers: forward; 5’-<u>GAATTC</u>ACCAGACAAGATGCCGAAGAGG-3’; reverse: 5’-<u>TCTAGA</u>GGTCAGTTCAGCAAGCACGACTC-3’, and the correct *Pacrg* cDNA was ligated into the pCS2+MT vector to create the PACRG/Myc plasmid. The PACRG /Myc and DNALI1/Flag plasmids were co-transfected into COS-1 cells by sing Lipofectamine™ 2000 transfection reagent (Invitrogen). After 48 h transfection, the cells were processed for co-immunoprecipitation assay. For co-immunoprecipitation assays, the cells were lysed with IP buffer (Beyotime, Jiangsu, China; Cat No. P0013) for 5 min and centrifuged at 10,000 g for 3∼5 min. The supernatant was pre-cleaned with protein A beads at 4°C for 30 min, and the pre-cleared lysate was then incubated with anti-MYC antibodies at 4°C for 2 h. The mixture was then incubated with protein A beads at 4°C overnight. The beads were washed with IP buffer three times and then re-suspended in 2x Laemmli sample buffer and heated at 95°C for 5 min. The samples were centrifuged at 3,000 g for 30 s, and the supernatant was then subjected to Western blot analysis with MYC and Flag antibodies.

### Gel filtration experiments

Full-length mouse *Pacrg* cDNA (amplified by a forward primer: 5’-<u>GAATTC</u>GGTGCCGCGCGGCAGCATGCCGAAGAGGACTAAACTG-3’ and a reverse primer: 5’-<u>GTCGAC</u>TCAGTTCAGCAAGCACGACTC-3’) was cloned into the upstream multiple clone site of the dual expression vector pCDFDuet-1 to create the PACRG/pCDFDuet-1 plasmid, and the translated protein was tagged with hexahistidine. The full-length mouse *Dnali1* cDNA (amplified by a forward primer: 5’-<u>GATATC</u>GATGATACCCCCAGCAGACTCTC-3’ and a reverse primer: 5’-<u>CTCGAG</u>TCACTTCTTCGGTGCGATAATG-3’) was inserted into the downstream multiple cloning site to create the PACRG/DNALI1/pCDFDuet-1 plasmid. The dual expression plasmid was transformed into the Rosetta II (DE3) (Invitrogen) E. coli strain, grown in Luria Bertani medium, and induced with 1 mM isopropyl-β-d-thiogalactopyranoside at an A600 ∼ 0.8 for 2 hours. The bacterial pellets from 1 L of growth media were resuspended in 30 mL of B-PER reagent (Thermo Scientific, Waltham, MA), and expressed proteins were purified from the lysis supernatant by Nickel affinity and gel filtration (Superdex 75 26/60, GE Healthcare, Chicago, IL) chromatography. The purified protein was concentrated and passed over the gel filtration column a second time before Western blot analysis.

### Luciferase complementation assay

HEK293 cells were grown in 12-well plates and transfected in triplicate with N-and C-luciferase fragments (supplied by Dr. James G Granneman, Wayne State University) fused to mouse PACRG and DNALI1 along with various controls as specified in the figure legends. The following primers were used to create the constructs: PACRG/C-Luc forward: 5’-The following primers were used to create the constructs: PACRG/C-Luc forward: 5’-<u>AAGCTT</u>CGATGGTGAAGCTAGCTGCCAAATG-3’, PACRG/C-Luc reverse: 5’-<u>ACCGGT</u>GGCACCAGGGTATGGAATATGTCCAC-3’, N-Luc/DNALI1 forward: 5’-<u>AAGCTT</u>ATGGCAGAGTTGGGCCTAAATGAG-3’, N-Luc/DNALI1 reverse: 5’-<u>ACCGGT</u>GGATCTTCAGATTCATATTTTGCCAG-3’. After transfection, cells were cultured for 24 h. Luciferase activities were measured as described previously (31) and readings were recorded using Veritas microplate luminometer. Experiments were performed three times independently, and the results are presented with standard errors.

### Western blot analysis

Tissue samples were collected from 3-4-month-old mice, and tissue extracts were obtained after lysis with buffer containing 50 mM Tris–HCl pH 8.0, 170 mM NaCl, 1% NP40, 5 mM EDTA, 1 mM DTT and protease inhibitors (Complete mini; Roche diagnostics GmbH, Basel, Switzerland). Protein concentration for lysates was determined using BCA reagent (Sigma-Aldrich, St. Louis, MO), and equal amounts of protein (50 μg/lane) were heated to 95°C for 10 min in sample buffer, loaded onto 10% sodium dodecyl sulfate-polyacrylamide gels, separated with electrophoresis, and transferred to polyvinylidene difluoride membranes (Millipore Corporation, Bedford, MA). Membranes were blocked (Tris-buffered saline solution containing 5% nonfat dry milk and 0.05% Tween 20 [TBST]) and then incubated with the indicated primary antibodies at 4°C overnight. After being washed in TBST, the blots were incubated with immunoglobulin conjugated to horseradish peroxidase for 1 h at room temperature. After washing, the target proteins were detected with Super Signal chemiluminescent substrate (Pierce, Thermo Scientific). The following primary antibodies were used: anti-His (1: 2000, Cat No: 70796-4, Novagen, Madison, WI); anti-DNALI1 (1: 10,000 for Proteintech 17601-1-AP, Rosemont, IL; and 1:2000 for Abcam, Cambridge, UK, Cat No: ab87075. This antibody has been discontinued); anti-GFP (1: 1000, Cat No: 11814460001, Roche); anti-MYC (1: 2000, Cat No: C-19 Sc788, Santa Cruz, Dallas, TX); and β-ACTIN (1: 2000, Cat No: 4967S, Cell Signaling Technology, Danvers, MA). Secondary antibodies include anti-Rabbit IgG (1: 2,000, Cat No: 711166152, Jackson ImmunoResearch, West Grove, PA) and anti-Mouse IgG (1: 2,000, Cat No: DI-2488, Vector Laboratories, Burlingame, CA). Each Western blot was performed using three independent biological replicates.

### Preparation of testicular cells and immunofluorescence analysis

Testes were separated from 3-4-month-old mice and incubated in a 15mL centrifuge tube with 5mL DMEM containing 0.5 mg/mL collagenase IV and 1.0 μg/mL DNase I (Sigma-Aldrich) for 30 min at 32^◦^C and shaken gently. Then the testes were washed once with PBS after centrifugation at 1,000 rpm for 5 min under 4^◦^C, and the supernatant was discarded. Afterwards, the cell pellet was fixed with 5 mL paraformaldehyde containing 0.1 M sucrose and shaken gently for 15 min at room temperature. After washing three times with PBS, the cell pellet was re-suspended with 2 mL PBS and loaded on positively charged slides. The slides were stored in a wet box at room temperature after air drying. Then, the spermatogenic cells were permeabilized with 0.1% Triton X-100 (Sigma-Aldrich) for 5 min at 37◦C. Finally, the samples were incubated overnight with primary antibodies (Anti-DNALI1: 1: 100, Abcam, Cat No: ab87075; Anti-PACRG: 1: 200, generated by our laboratory; Anti-α-tubulin, 1:200, Sigma, Cat No: T9026-2mL; Anti-MEIG1, 1:400, generated by our laboratory; Anti-SPAG16L, 1: 200, generated by our laboratory; anti-DYNC1H1: 1: 100, Santa Cruz Biotechnology, Cat No: sc-7527). After washing three times with PBS, the slides were incubated with secondary antibodies for 1h at room temperature. CyTM3 AffiniPure F (ab’)2 Fragment Donkey Anti-Rabbit IgG (H + L) (1: 200, Jackson ImmunoResearch, Cat No: 711166152) and DyLight 488 Horse Anti-Mouse IgG Antibody (1: 400, Vector Laboratories, Cat No: DI-2488) are two secondary antibodies used in this experiment. Images were captured using multiple microscopes in separate locations: confocal laser scanning microscopy (Zeiss LSM 700, Virginia Commonwealth University), Olympus IX-81 microscope equipped with a spinning-disc confocal unit (Dr. James G. Granneman, Wayne State University), and Nikon DS-Fi2 Eclipse 90i Motorized Upright Fluorescence Microscope (The C.S. Mott Center for Human Growth and Development, Department of Obstetrics & Gynecology, Wayne State University).

### Generation of *Dnali1* conditional knockout (cKO) mice

*Dnali1^flox/flox^* mice were generated at the Center for Mouse Genome Modification at University of Connecticut, and *Stra8-iCre* mice were purchased from the Jackson Laboratory (Stock No: 008208). Cre recombinase was shown to be only active in male germ cells (32). To generate the germ cell-specific *Dnali1* cKO mouse model, 3-4-month-old Stra8-cre males were crossed with 3-4-month-old *Dnali1^flox/flox^* females to obtain *Stra8-iCre*; *Dnali1^flox/+^* mice. The 3-4-month-old *Stra8-iCre*; *Dnali1^flox/+^* males were crossed back with 3-4-month-old *Dnali1^flox/flox^* females again. The *Stra8-iCre*; *Dnali1^flox/flox^* were considered to be the homozygous cKO mice, and *Stra8-iCre*; *Dnali1^flox/+^* mice were used as the controls. Genomic DNA was isolated to genotype the offspring. The following primers were used for genotyping: *Stra8-iCre* forward: 5′-GTGCAAGCTGAACAACAGGA-3′; *Stra8-iCre* reverse: 5′-AGGGACACAGCATTGGAGTC-3′. An 844 bp PCR product is amplified from the floxed allele of the *Dnali1* with the following primers: forward: 5′-CCTGTGGGAAAGCTAACCCAGC-3’(DliScF5), and reverse: 5′-GCTGGGGATGCGGTGGGCTC-3′ (BGHpAr). A 129 bp PCR product is amplified from the wild-type allele using the following primers: forward: 5’-GACAGGGATGGAGGTTGGGAG-3’, and reverse: 5’-GAATGAGTGGTCAGGCCTCTG-3’. The *Pacrg* mutant mice were purchased from Jackson Laboratory (stock number: 000567, 27).

### Assessment of male fertility and fecundity

Sexually mature *Dnali1* cKO and control male mice were each mated with a 2-4-month old wild-type female for at least 2 months. The presence of vaginal plugs was checked, and the pregnancy of females was recorded. The number of pups was counted the day after birth. Average litter sizes are presented as the number of total pups born divided by the number of mating cages.

### Sperm parameters

After breeding studies, the male mice were euthanized by cervical dislocation following anesthesia. Sperm were collected after swimming out from the cauda epididymis in 37°C PBS. Cells were counted using a hemocytometer chamber under a light microscope, and sperm number was calculated by standard methods. Motility percentages and velocities (average path velocity) were then analyzed using Image J (National Institutes of Health, Bethesda, MD) and the plug-in MTrackJ.

### HE staining of testis and epididymis sections

Testes and epididymides of adult mice were collected and fixed in 4% paraformaldehyde (PFA) in phosphate-buffered saline (PBS) at 4°C overnight. The tissues were embedded in paraffin, sectioned at 5μm thickness, deparaffined, and stained with hematoxylin and eosin using standard procedures. Slides were examined using a BX51 Olympus microscope (Olympus Corp., Melville, NY, Center Valley, PA), and photographs were taken with a ProgRes C14 camera (Jenoptik Laser, Germany).

### Transmission electron microscopy

Mouse testes and sperm from adult control and *Dnali1* cKO mice were collected after swimming out from the cauda epididymis in 37°C PBS solution followed by centrifugation at 1,000 × g for 10 min under 4°C and re-suspended in fixation buffer (0.2M HEPES, 8mM CaCl_2_, 8% PFA, 10% Glutaraldehyde, PH7.4). Electron microscopy was performed using the method described previously (33).

### Statistical analysis

Statistical analyses were performed using Student’s *t* test. *p < 0.05 was considered as significant. Graphs were created using Microsoft Excel.

## Results

### DNALI1 is a binding partner of PACRG

PACRG, a major spermatogenesis regulator, was used as bait for a yeast two-hybrid screen. The sequencing results of positive clones revealed that MEIG1 was the major binding partner as previously described (26). In addition, DNALI1 encoding clones were identified multiple times (Supplemental Table 1). A direct yeast two-hybrid assay was conducted to confirm the interaction between PACRG and DNALI1. Like the positive control, the yeast co-transformed with the two plasmids expressing DNALI1 and PACRG grew on the selection medium (Figure 1A), indicating that the two proteins interact in yeast. To further examine interaction of the two proteins, CHO cells were transfected with the plasmids expressing the two proteins. When the CHO cells expressed PACRG/GFP only, the protein was present in the cytoplasm (Figure 1B, a) while DNALI1/FLAG was present as a vesicle located on one side of the nucleus (Figure 1B, b). When CHO cells expressed both proteins, PACRG/GFP was present as a vesicle and was co-localized with DNALI1/FLAG (Figure 1Bc-e). In addition, we transfected COS-1 cells with DNALI1/FLAG and PACRG/Myc expression plasmids and conducted a co immunoprecipitation assay. The MYC antibody pulled down both the Myc-tagged 28 kDa PACRG and Flag-tagged 34 kDa DNALI1 proteins, suggesting an interaction between these proteins (Figure 1C). Finally, interaction between DNALI1 and PACRG was examined by luciferase complementation assay. We observed that HEK cells co-expressing PACRG/C-Luc and N-Luc/DNALI1 showed robust luciferase activity, while the cells expressing either PACRG/C-Luc or N-Luc/DNALI1 only showed baseline activity (Figure 1D).

**Figure 1.**
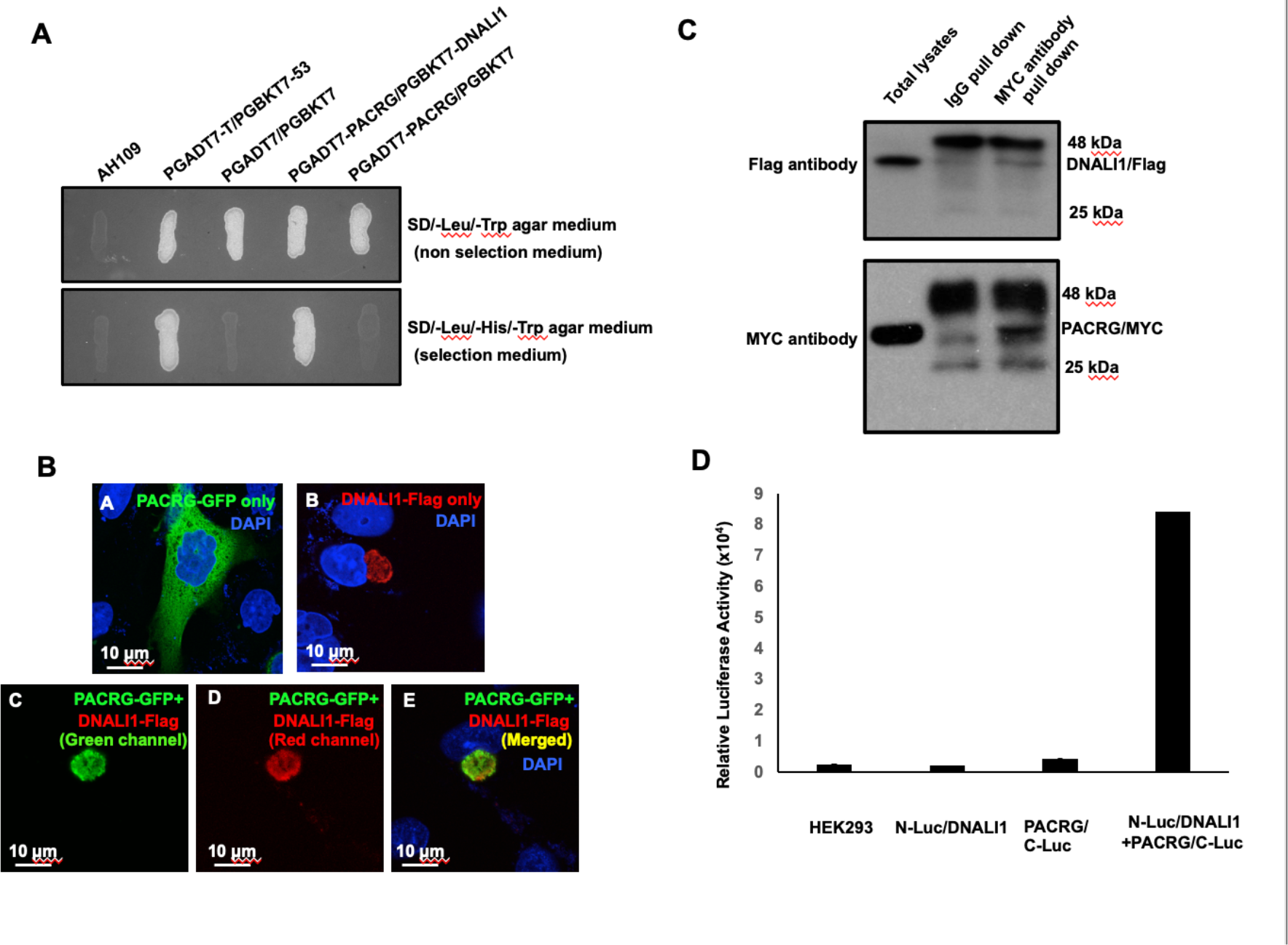
DNALI1 associates with PACRG, a major spermatogenesis regulator. A. Direct yeast two-hybrid assay to examine the interaction between PACRG and DNALI1. Pairs of indicated plasmids were co-transformed into AH109 yeast, and the transformed yeast were grown on either selection plates (lacking leucine, histidine and tryptophan) or non-selection plates (lacking leucine and tryptophan). Notice that all the yeast except AH109 grew on the non-selection plate. Yeast expressing PACRG/DNALI1 and P53/large T antigen pairs grew on selection plate. B. DNALI1 co-localizes with PACRG in CHO cells. When expressed alone, PACRG/GFP was present in the cytoplasm, and DNALI1/FLAG as a vesicle located closed to the nucleus. When the two proteins were co-expressed, PACRG/GFP was present as a vesicle and co-localized with DNALI1/FLAG. (Images taken with confocal laser scanning microscopy (Zeiss LSM 700), Virginia Commonwealth University) C. Co-immunoprecipitation of DNALI1/FLAG with PACRG/Myc. COS-1 cells were transfected with plasmids to co-express DNALI1/FLAG and PACRG /Myc. The cell lysate was immunoprecipitated with anti-MYC antibody and then analyzed by Western blotting with anti-MYC and anti-FLAG antibodies. The cell lysate immunoprecipitated with a mouse normal IgG was used as a control. The anti-MYC antibody pulled down both PACRG/MYC and DNALI1/FLAG. D. Interaction of PACRG with DNALI1 in HEK293 cells as determined by G. princeps luciferase complementation assay. HEK293 cells were transfected with the indicated plasmids, and luciferase activity was evaluated 24 h after transfection. The cells expressing both N-Luc/DNALI1 and PACRG/C-Luc reconstituted activity.

### DNALI1 is co-purified with His-tag PACRG when the two proteins are co-expressed in BL21 bacteria

In order to examine the association between DNALI1 and PACRG in bacteria, we co-expressed DNALI1 and His-tagged PACRG in BL21 cells, and a gel filtration experiment was conducted to purify the His-tagged PACRG protein into continuous fractions. Detection of His-tagged PACRG and DNALI1 proteins in the eluted fractions was examined by Western blot using specific antibodies. In line with the above interaction data, DNALI1 protein was co-purified with PACRG protein in the same fractions (Figure 2).

**Figure 2.**
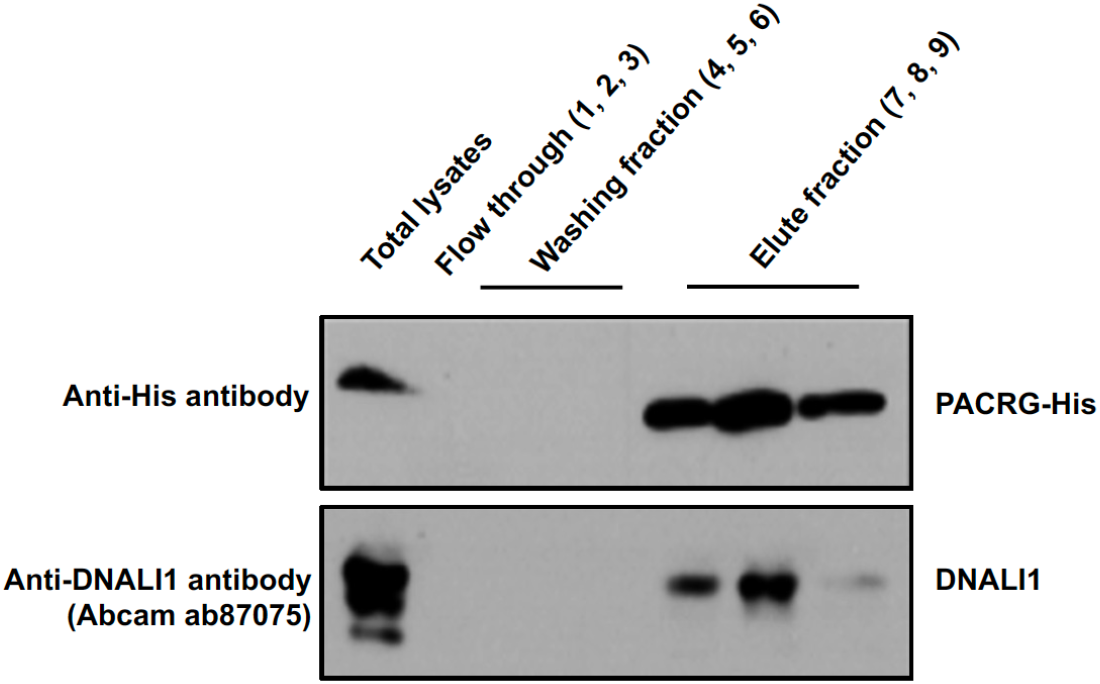
Co-purification of DNALI1 with His-tagged PACRG from bacteria lysates expressing the two proteins. His-tagged PACRG and DNALI1 were co-expressed in BL21 bacteria and His-tagged PACRG was purified from the bacteria lysates by gel filtration experiments, and presence of the His-tagged PACRG and non-tagged DNALI1 in the fractions were examined by western blot analysis. Notice that the DNALI1 protein was present in the same fractions of His-tagged PACRG, indicating that the two proteins associate in the bacteria lysates.

### DNALI1 stabilizes PACRG in mammalian cells and bacteria

Functional association between PACRG and DNALI1 was further supported by the fact that DNALI1 stabilized PACRG in mammalian cells and bacteria. PACRG alone was not stable in the transfected COS-1 cells with very low protein amount detected (27). Co-expression of DNALI1 dramatically increased PACRG level (Figure 3A). In BL21 bacteria, no PACRG protein was detectable when the less sensitive Pico system was used for Western blot analysis when the bacteria were transformed with PACRG/pCDFDuet-1 plasmid and induced by IPTG. However, the PACRG protein was indeed expressed as it was detectable when the higher sensitive Femto system was used for Western blot analysis. When the BL21 bacteria were transformed by PACRG/DNALI1/pCDFDuet-1 plasmid so that both PACRG and DNALI1 were expressed, the PACRG was easily detectable with the Pico system (Figure 3B), indicating that wild-type DNALI1 also stabilized PACRG protein in bacteria.

**Figure 3.**
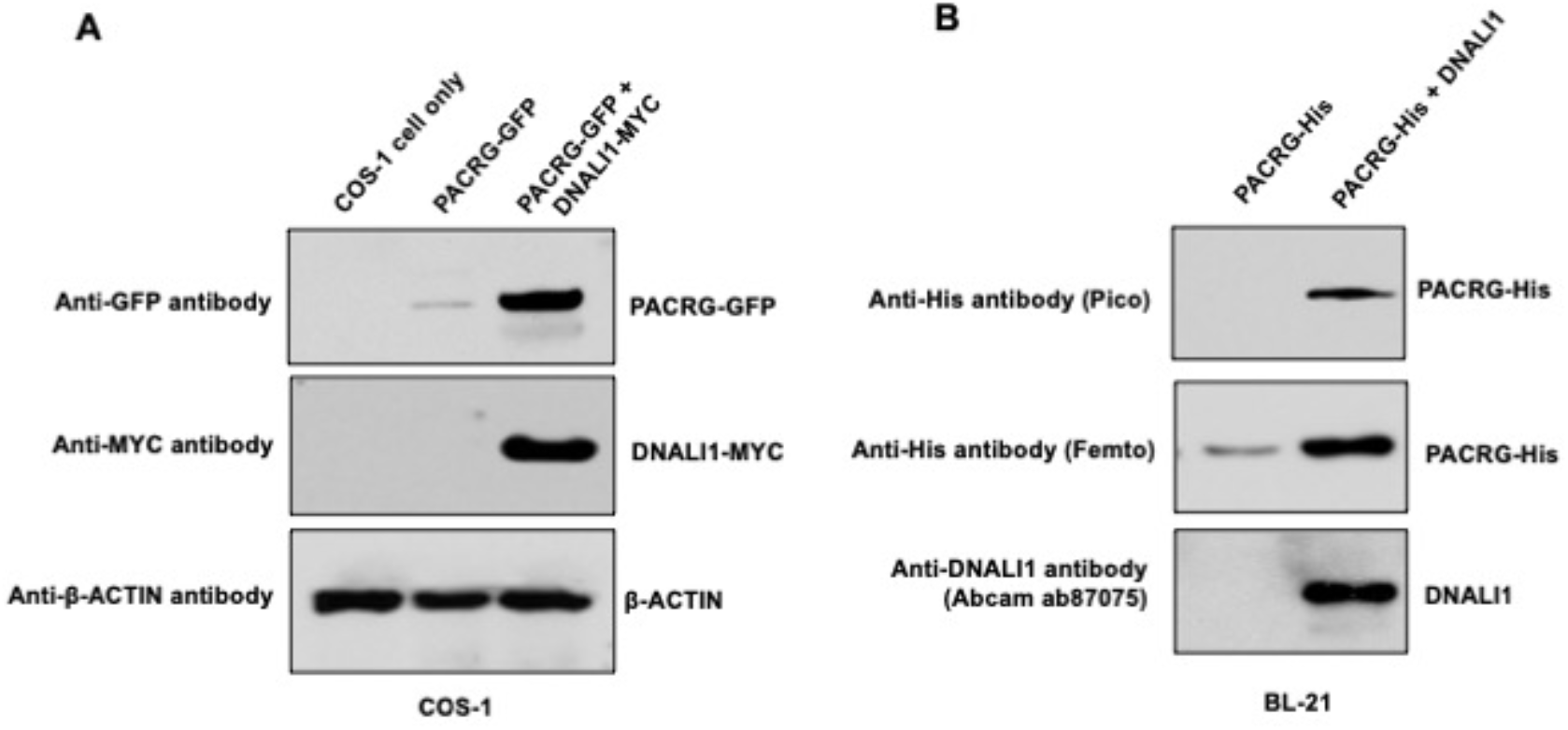
DNALI11 stabilizes PACRG in mammalian cells and bacteria. A. DNALI1 stabilizes PACRG in COS-1 cells. Mouse PACRG-GFP expression is increased when DNALI1 is co-expressed in transfected COS-1 cells in transient expression experiment. B. DNALI1 stabilizes PACRG in bacteria. Notice that PACRG was only detectable by the high sensitivity Femto system in Western blot analysis when the bacteria were transformed with PACRG/pCDFDuet-1 plasmid. However, when the bacteria were transformed with PACRG/DNALI1/pCDFDuet-1 plasmid to express DNALI1 protein, PACRG was also detectable by less sensitive Pico system.

### Localization of DNALI1 in the manchette is not dependent on PACRG

DNALI1 localization in germ cells was examined by immunofluorescence staining. In elongating spermatids, DNALI1 co-localized with the α-tubulin, a manchette marker. Double staining with an anti-DNALI1 polyclonal antibody and an anti-PACRG monoclonal antibody revealed that the two proteins also co-localized (Figure 4). In *Pacrg* mutant mice, DNALI1 was still present in the manchette, indicating that the localization of DNALI1 in the manchette is not dependent on PACRG (Supplemental Figure 1).

**Figure 4.**
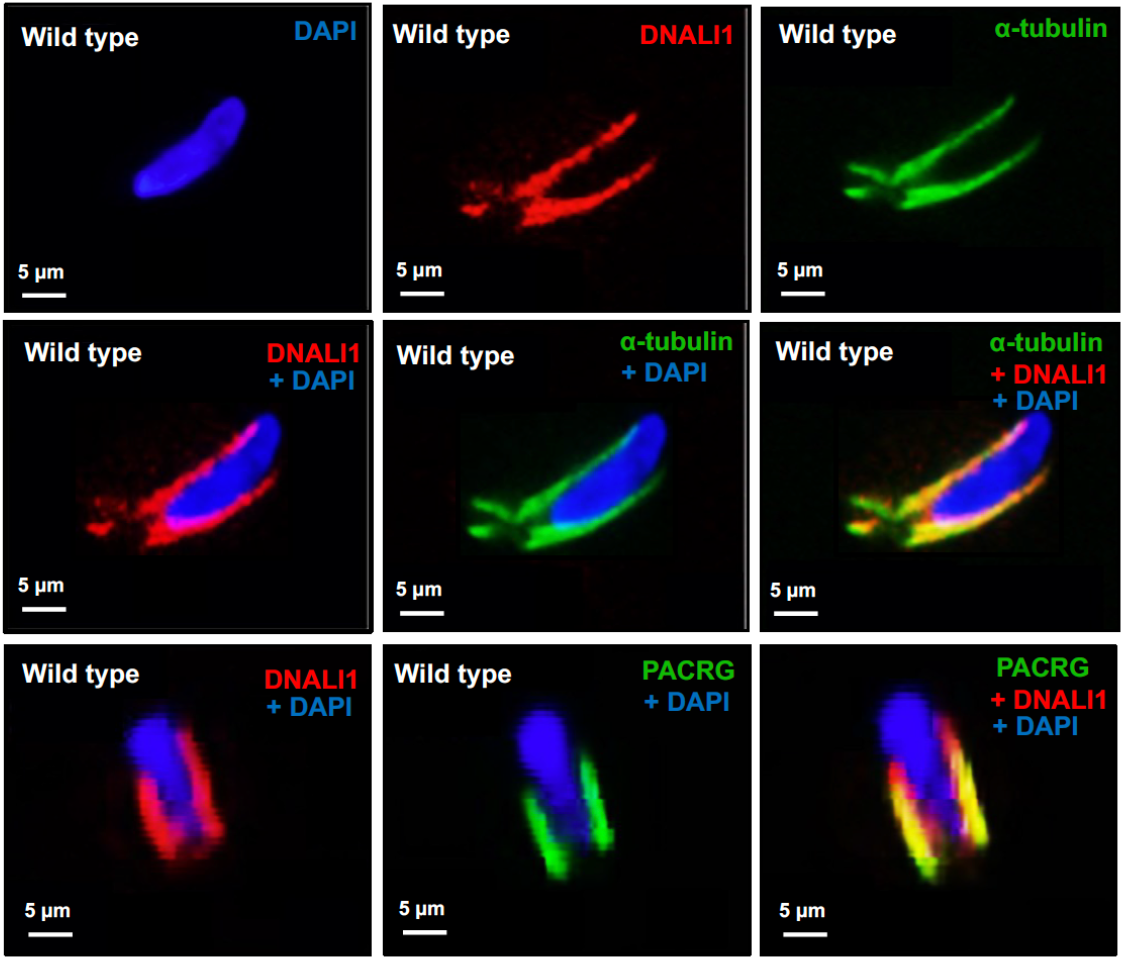
Localization of DNALI1 in male germ cells of wild-type mice. Localization of DNALI1 in isolated germ cells was examined by immunofluorescence staining. DNALI1 protein localizes with α-tubulin, a manchette marker in elongating spermatids (top and middle panels). PACRG also co-localized with DNALI1 in the elongating spermatids (bottom panel). DNALI1 seems to be closer to the nuclear membrane, and PACRG is on the surface of DNALI1. (Images taken with confocal laser scanning microscopy (Zeiss LSM 700), Virginia Commonwealth University)

### Inactivation of mouse *Dnali1* gene resulted in infertility associated with significantly reduced sperm number, motility and increased abnormal

To explore the role of DNALI1 in spermatogenesis and male fertility, a floxed *Dnali1* line was generated and the floxed *Dnali1* mice were crossed with *Stra8-Cre* mice so that the *Dnali1* gene was specifically disrupted in male germ cells (*Dnali1* cKO). Genotyping using specific primer sets indicated that homozygous mutant mice were obtained (Supplemental Figure 2). Examination of total testicular DNALI1 protein expression by Western blot revealed that DNALI1 was almost absent in the conditional knockout mice (Figure 5A). Homozygous mutant mice did not show any gross abnormalities. To test fertility of these mutant mice, 2-3-month-old controls and homozygous mutant males were bred with 2-3-month-old wild-type females for 2 months. All the control mice, including wild-type and heterozygous males, were fertile. All homozygous mutant males examined were infertile (Figure 5B). Epididymal sperm number, motility and morphology from the control and homozygous mutant mice were examined. The sperm count and motility was dramatically reduced in the mutant mice (Figure 5C, D, E **and** Supplemental Figure 3). Sperm from the control mice showed normal morphology (Figure 6A). Multiple abnormalities in sperm were observed in the mutant mice, including head defects, uneven thickness of tails and sperm bundles lacking separation (also called individualization or disengagement) (Figure 6B-F, Supplemental Figure 4).

**Figure 5.**
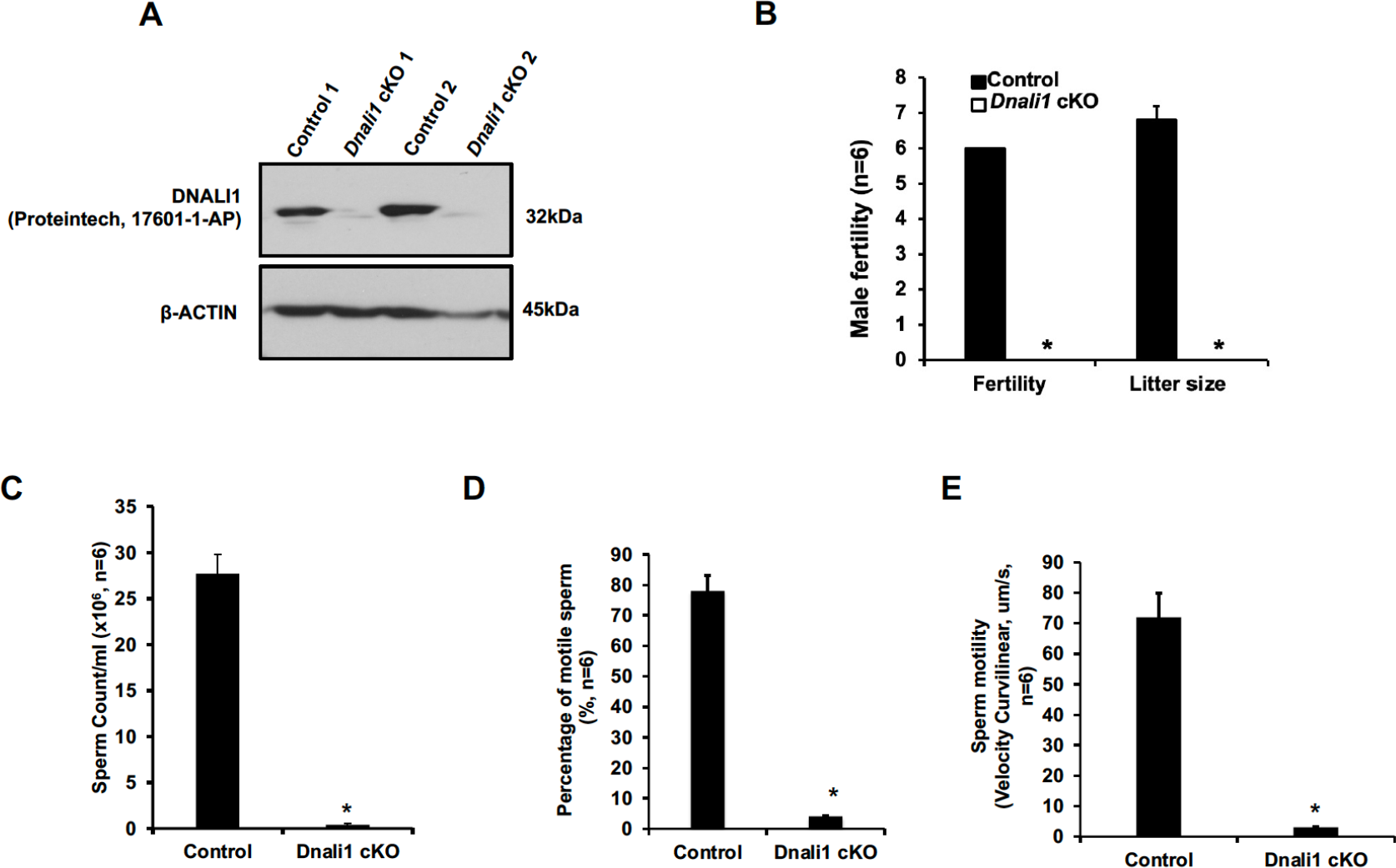
Male germ cell-specific *Dnali1* knockout (*Dnali1* cKO) mice were infertile associated with significantly reduced sperm number and motility. A. Representative Western blot result showing that DNALI1 protein was almost absent in the testis of *Dnali1* cKO mice. B. Male fertility of control and *Dnali1* mutant mice. Six controls and six *Dnali1* cKO mice were examined. Fertility and litter size were recorded for each mating. Notice that all mutant males were infertile. C. Sperm count was significantly reduced in the *Dnali1* cKO mice (n=6). D. Percentage of motile sperm in the control and *Dnali1* cKO mice (n=6). E. Sperm motility was significantly reduced in the *Dnali1* cKO mice (n=6). Statistically significant differences: *p < 0.05.

**Figure 6.**
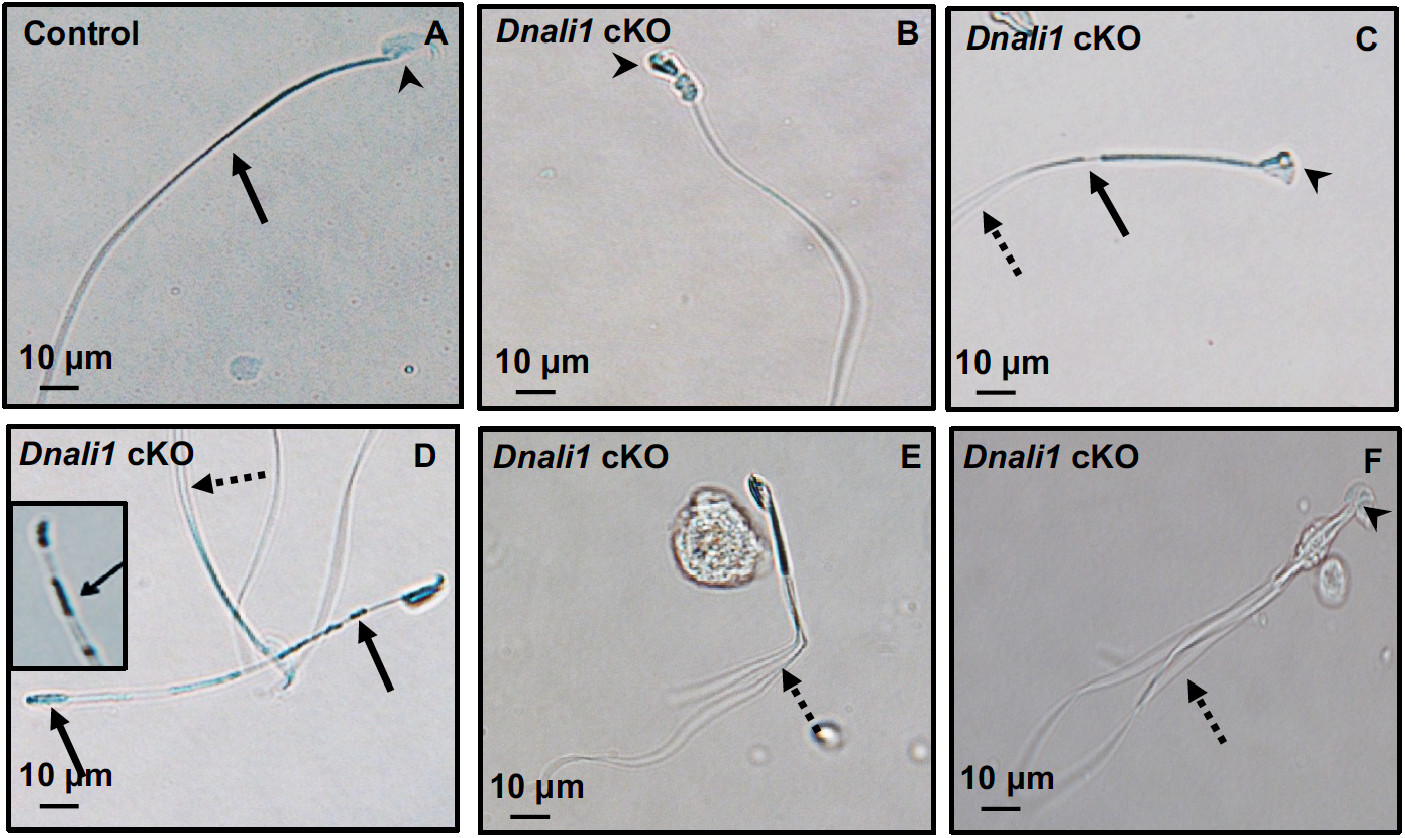
Abnormal sperm morphologies in the *Dnali1* cKO mice. Representative epididymal sperm of control (A) and *Dnali1* cKO mice (B-F) examined by DIC microscopy. Sperm in the control mice showed normal head (A, arrowhead) and flagella (A, arrow). Multiple abnormalities were observed in the *Dnali1* cKO mice, including distorted heads (B, C and F, arrowheads), uneven thickness tails (C, D, arrows). Some sperm showed multiple flagella (C, D, E, F, dashed arrows).

### Spermatogenesis was affected in the *Dnali1* cKO mice

Reduced epididymal sperm number and aberrant sperm morphology in *Dnali1* cKO mice suggests impaired spermatogenesis. Therefore, histology of the testes and epididymides in adult control and *Dnali1* mutant mice was examined. In control mice, the seminiferous tubule epithelium showed elongated spermatid heads embedded as bundles among the round spermatids and developing tails extending into the lumen in stages I-III (Figure 7Aa). *Dnali1* cKO mice exhibited abnormal elongating spermatid heads with compressed shapes and failure to form elongated bundles. Round spermatids were seen sloughing abnormally into the lumen in stages I-III (Figure 7Ab **and** c). In stage XII, the control seminiferous tubule epithelium showed step 12 elongating spermatid bundles with large pachytene spermatocytes in meiotic division (Figure 7Ad). However, in the *Dnali1* cKO mice, stage XII showed abnormal elongating spermatid heads without normal bundle formation. Additionally, the phagocytosis of thin heads of step 16 spermatids was present in the epithelium, which is evidence of spermiation failure (Figure 7Ae **and** f). Numerous mature sperm were concentrated in the cauda epididymis of control mice (Figure 7B, **left**). In *Dnali1* cKO mice, the cauda epididymal lumen contained fewer sperm, along with abnormal sperm heads and tails and sloughed round spermatids (Figure 7B, **right**).

**Figure 7.**
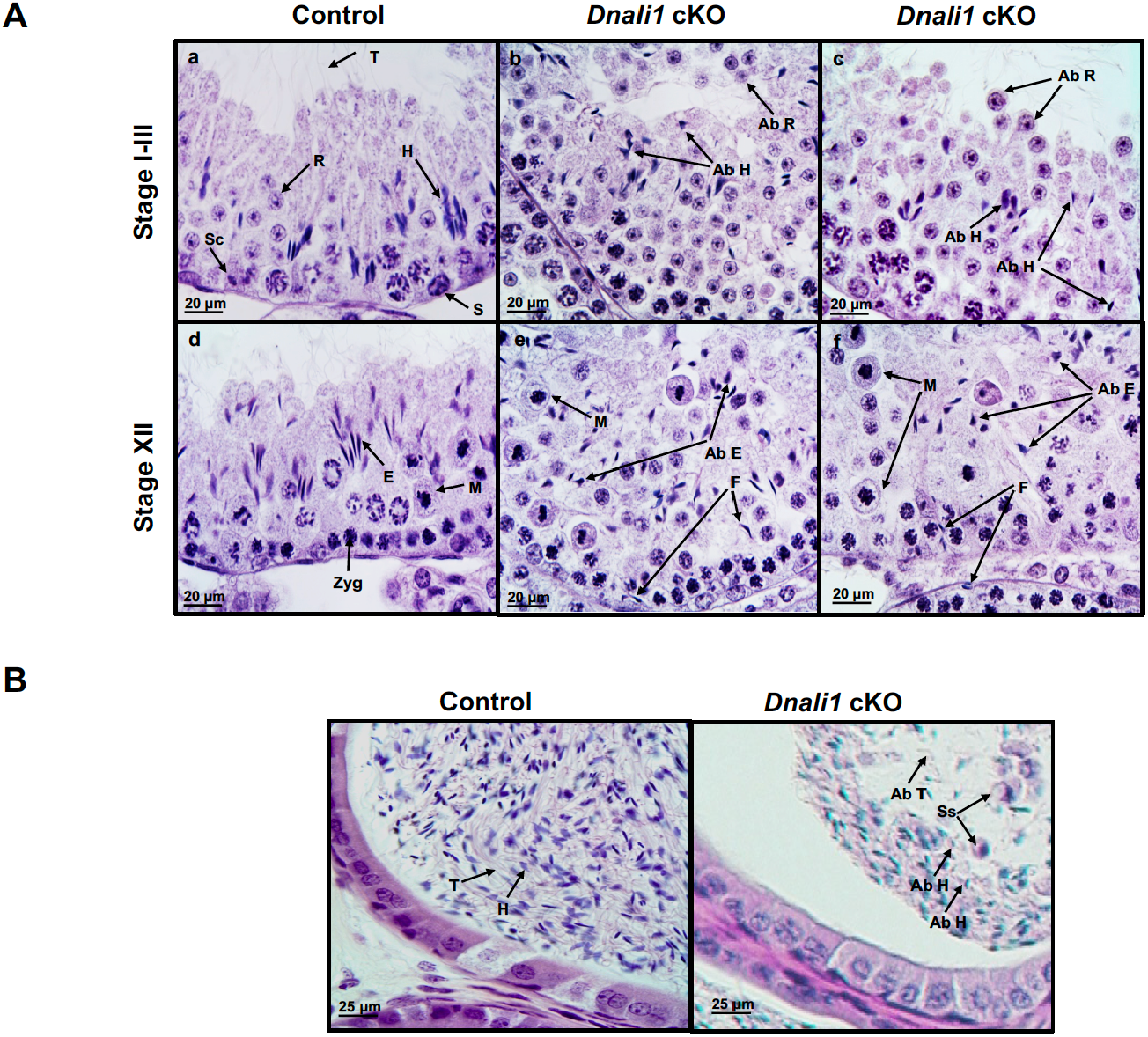
Histology analysis for the adult c ontrol and *Dnali1* cKO mice. A. Histological evaluation of testes from control and *Dnali1* cKO mice, with selected images from Stages I-III (a-c) and Stages XII (d-f). a) Control testis seminiferous tubule epithelium showing elongated spermatid heads (H) embedded as bundles among the round spermatids (R) and developing tails (T) extending into the lumen. Sc, Sertoli cell; S, spermatogonia. b-c) *Dnali1* cKO with abnormal elongating spermatid heads (Ab H) with compressed shapes and failure to form elongated bundles. Round spermatids are seen sloughing abnormally into the lumen (Ab R). d) Control showing step 12 elongating spermatid bundles (E) with large pachytene spermatocytes in meiotic division (M). Zygotene spermatocyte, Zyg. e-f) *Dnali1* cKO with abnormal elongating spermatid heads (Ab E) without normal bundle formation. There is evidence of failure of spermiation (F), as phagocytosis of thin heads of step 16 spermatids are also present in the epithelium. Normal meiotic figures (M) are present in Stage XII. B. Representative histology of epididymis. The control cauda epididymis shows the lumen filled with normal sperm heads (H) and tails (T). In the *Dnali1* cKO male, the cauda epididymal lumen contains numerous sperm with abnormal heads (Ab H) and tails (Ab T) and sloughed round spermatids (Ss).

### Ultrastructural changes in the seminiferous tubules of the *Dnali1* cKO mice

To investigate the structural basis for the molecular changes observed in the absence of DNALI1, the seminiferous tubule and cauda epididymal sperm ultrastructure was examined. In control mice, the nuclei have normal elongated shapes with condensed chromatin (Figure 8A), and the flagellum contains a normal “9 + 2” axoneme structure surrounded by accessory structures, including the mitochondrial sheath (Figure 8B) and a fibrous sheath (Figure 8C). In *Dnali1* cKO mice, abnormal shapes of the nuclei were frequently seen (Figure 8D) and retained cytoplasmic components (not fully resorbed as part of the residual body) were present in the developed sperm (Figure 8E, Supplemental Figure 5A). Importantly, the two longitudinal columns were usually not associated with microtubule doublets 3 and 8, and the two semi-circumferential ribs showed defective or asymmetric formation in the principal piece of flagella; some sperm also had abnormal cell membranes (Figure 8F, Supplemental Figure 5B, C). Lastly, in some sperm the core “9+2” axoneme was incomplete or disorganized (Figure 8G-I).

**Figure 8.**
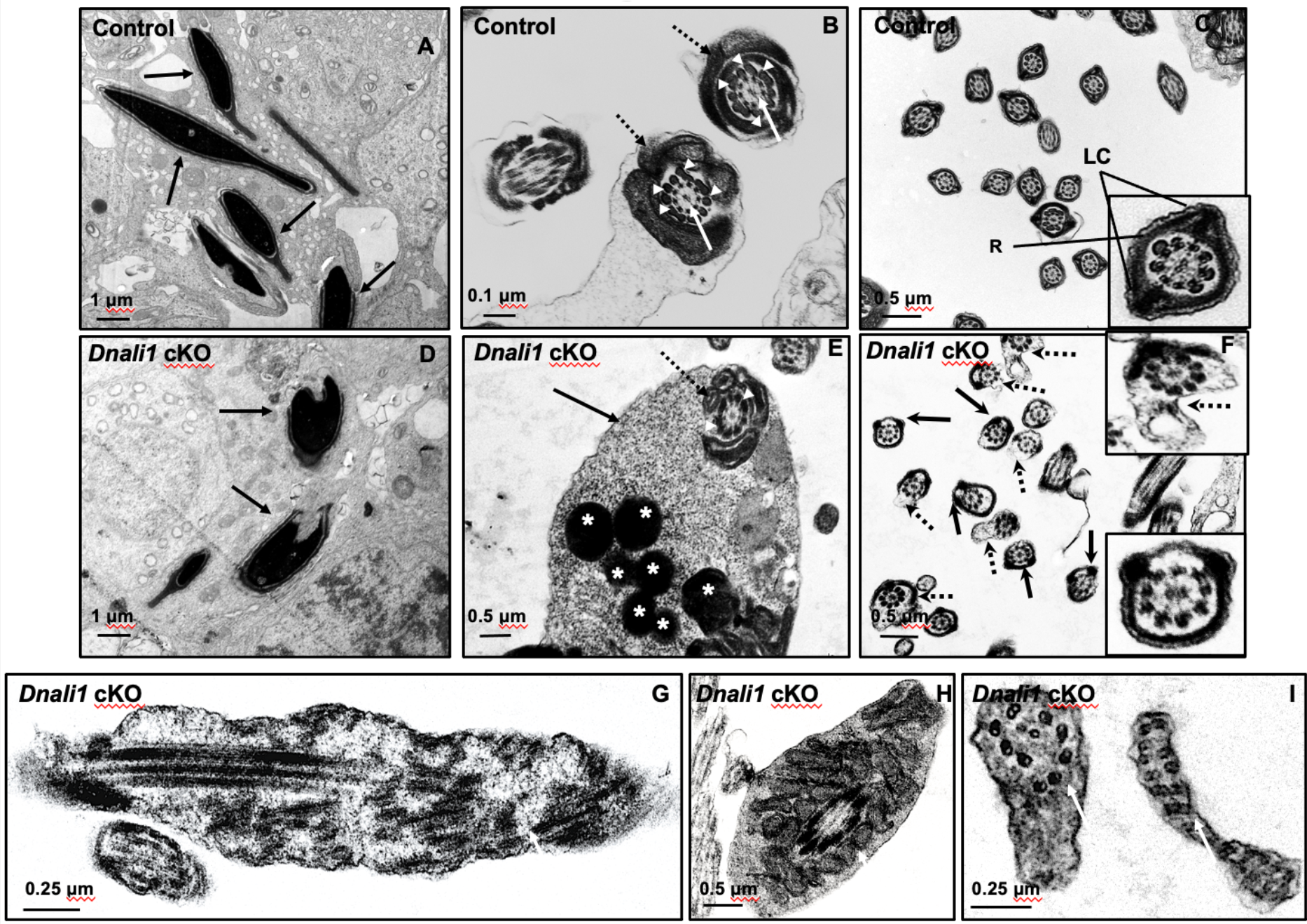
Ultra-structural changes of testicular sperm of control and *Dnali1* cKO mice. The ultrastructure of testicular sperm from the control (A, B and C) and *Dnali1* cKO (D-I) mice were analyzed by TEM. A. Control testis seminiferous tubule epithelium showing nuclei with normally condensed chromatin (arrows). B. Control mouse showed the normal middle piece of flagella with normal “9 + 2” axoneme structure in the center (white arrows) surrounded by mitochondrial sheath (black dotted arrows) and ODF (white arrow heads). C. Normal principal piece of flagella from a control mouse, which is characterized by the presence of a complete fibrous sheath surrounding the axoneme. The fibrous sheath consists of two longitudinal columns (LC) connected by semicircumferential ribs (R). The two longitudinal columns are associated with microtubule doublets 3 and 8 with the two semicircumferential ribs are symmetrical. D. Abnormally condensed chromatin in the *Dnali1* cKO mouse (black arrows). E. A representative image of the middle piece in a flagellum from a *Dnali1* cKO mouse. The ODF (white arrow heads) and mitochondrial sheath (black dotted arrow) were present, but cytoplasm residue (black arrow) remained with a number of the lysosomes inside (white stars). F. The flagella show disorganized fibrous sheath structure in the *Dnali1* cKO mouse. Noticed that the two longitudinal columns are not associated with microtubule doublets 3 and 8 in some flagella, and the two semicircumferential ribs showing defective or asymmetric organization (black arrows and the lower, right insert); some flagella also have disrupted membranes (dashed arrows and upper, right insert). G-I: the flagella show disrupted axoneme in the *Dnali1* cKO mice (white arrows).

### Inactivation of *Dnali1* in male germ cells did not change protein levels but changed the localization of the downstream proteins

To examine if the loss of DNALI1 in male germ cells affected protein levels of MEIG1, PACRG and SPAG16L, Western blotting was conducted using the testis lysates of control and *Dnali1* cKO mice. There was no significant difference in protein levels of MEIG1, PACRG and SPAG16L between the control and *Dnali1* cKO mice (Figure 9A). Localization of MEIG1 and SPAG16L was further examined. MEIG1 was present in the cell bodies of spermatocytes and round spermatids of control mice, and protein localization was not changed in the two cell types in *Dnali1* cKO mice (27, 29, Figure 9B). Similarly, SPAG16L was present in the cytoplasm of spermatocytes and round spermatids in both control and *Dnali1* cKO mice (27, 29, Figure 9C). In contrast, while MEIG1 and SPAG16L were present in elongating spermatids of *Dnali1* cKO mice, they did not co-localized in the manchette as observed for control mice (Figure 9D, Supplemental Figure 6).

**Figure 9.**
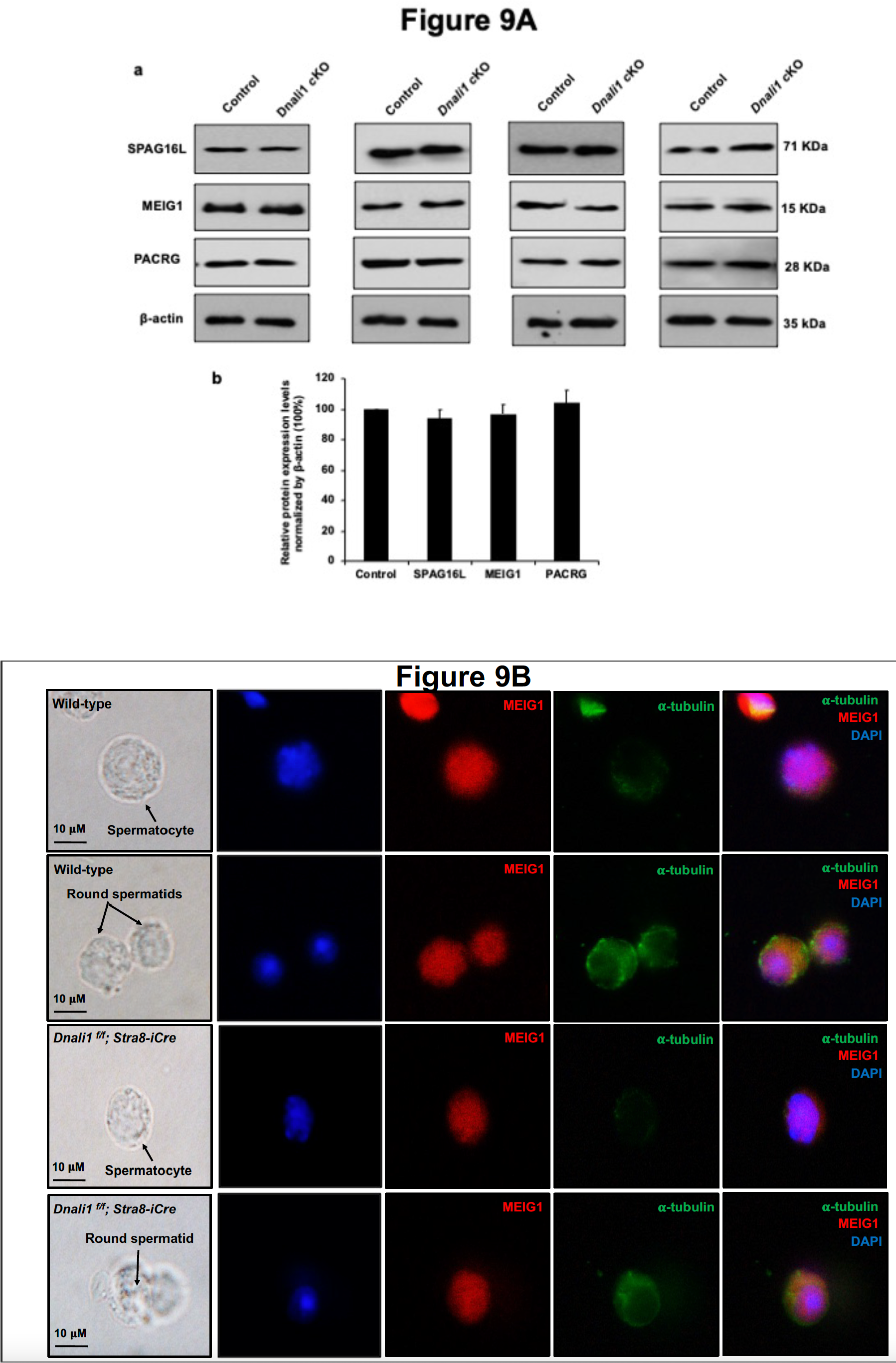
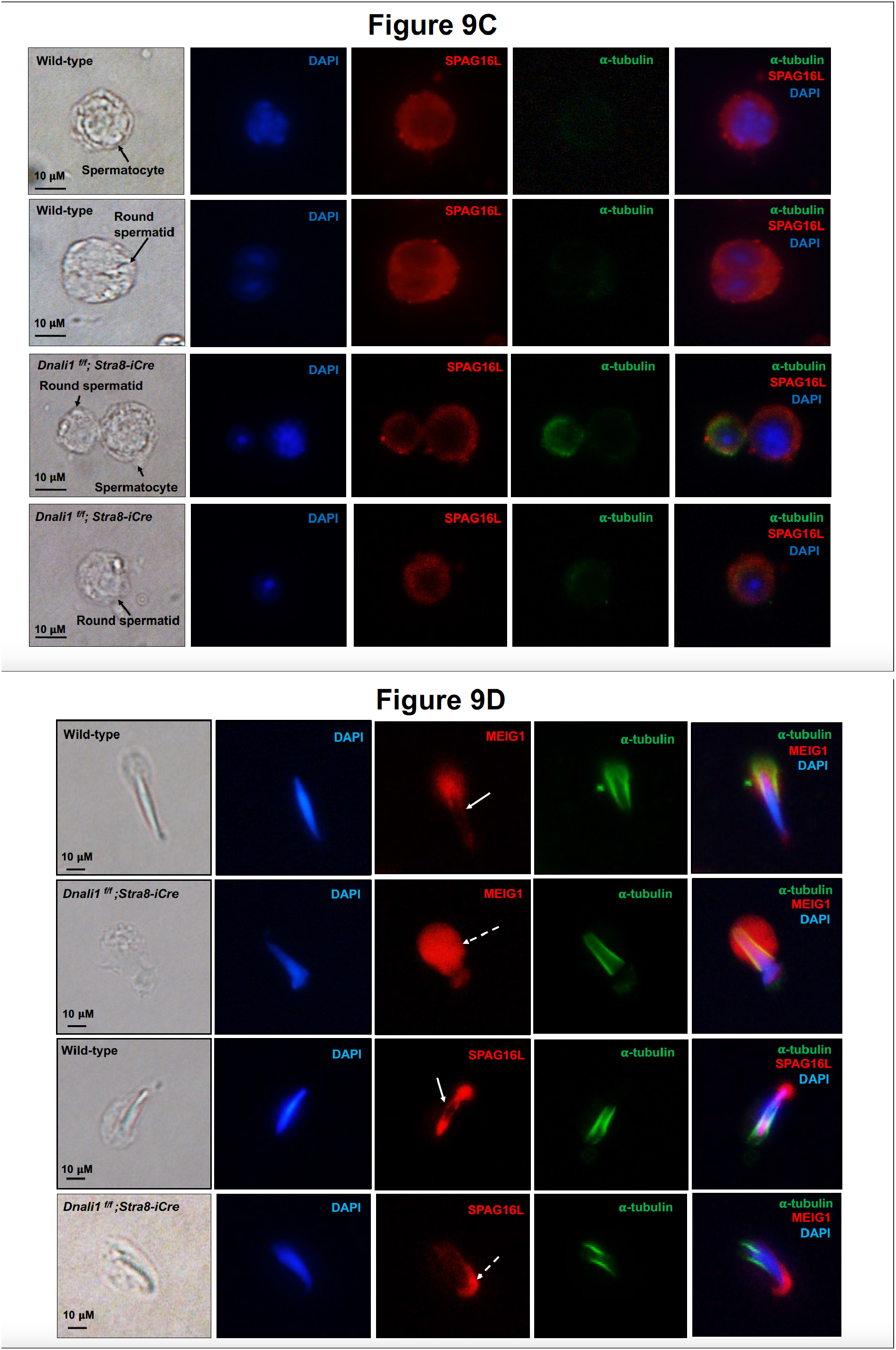
Expression levels and localization of DNALI1 downstream proteins in the *Dnali1* cKO mice. A. Analysis of testicular expression of MEIG1, PACRG and SPAG16L in control and *Dnali1* cKO mice by Western blot. Compared with control mice, there was no significant change in the expression level of these three proteins in the *Dnali1* cKO mice. a. Representative Western blot results; b. Statistical analysis of the protein levels normalized by β-actin. n=4. B. Localization of MEIG1 in spermatocytes and round spermatids of the control and *Dnali1* cKO mice by immunofluorescence staining. There is no difference between the control and the *Dnali1* cKO: MEIG1 is present in cell bodies in both genotypes. C. Localization of SPAG16L in spermatocytes and round spermatids of the control and *Dnali1* cKO mice by immunofluorescence staining. There is no difference between the control and the *Dnali1* cKO: SPAG16L is present in cell bodies in both genotypes. The white dashed arrow points to an elongating spermatid, and the SPAG16L is present in the manchette. D. Localization of MEIG1 and SPAG16L in elongating spermatids of the control and *Dnali1* cKO mice. Both MEIG1 and SPAG16L are present in the manchette in the control mice (white arrows); however, they are no longer present in the manchette in the *Dnali1* cKO mice (dashed white arrows). (Images taken with a Nikon DS-Fi2 Eclipse 90i Motorized Upright Fluorescence Microscope).

## Discussion

The *Parcg* gene is a reverse strand gene located upstream of the *Parkin* gene, involved in Parkinson’s disease (34). Genetic disruption of *Pacrg* in mouse phenocopies the infertility phenotype of *Meig1* knockout mice (27, 29, 35–38). Our previous studies demonstrated that MEIG1 and PACRG form a complex in the manchette for protein cargo transport to build the sperm flagellum. We further discovered that four amino acids on the same surface of MEIG1 protein are involved in its interaction with PACRG (28, 29). However, it was not clear how the MEIG1/PACRG complex associates with the manchette microtubules. Given that MEIG1 is a downstream binding partner of PACRG, and the fact that PACRG does not associate with microtubules directly, we hypothesized that there must be other upstream players involved in the manchette localization of the MEIG1/PACRG complex. Therefore, we performed a yeast two-hybrid screen using a PACRG component as bait. Besides MEIG1, DNALI1 was identified to be major binding partner. The interaction between PACRG and DNALI1 was further confirmed by other experiments described in this study, including the gel filtration and protein complementation experiments.

The functional assays reported here also support the observed interaction between PACRG and DNALI1. Previously, we discovered that mouse PACRG was not stable but could be stabilized by MEIG1 in both bacteria and mammalian cells (28). The studies here showed that DNALI1 also stabilized PACRG, suggesting a dual association of PACRG with MEIG1 and DNALI1. Similar to the MEIG1 protein, co-expression of DNALI1 also prevented degradation of PACRG in both bacteria and mammalian cells, which is consistent with the reported regulation of PACRG by the ubiquitin-proteasomal system (UPS) (39). It is possible that the binding site on the PACRG recognized by the UPS system is protected by DNALI1, providing PACRG protein more stability.

This association between PACRG and DNALI1 was also supported by *in vivo* studies. During the first wave of spermatogenesis, expression of the DNALI1 protein was dramatically increased during the spermiogenesis phase, which was similar to the PACRG protein (22, 28) and specific to the period of spermatid elongation and formation of flagella. However, prior to spermiogenesis, in late meiotic germ cells, the DNALI1 signal was weak (22). In elongating spermatids, DNALI1 was localized in the manchette, coinciding with the localization of PACRG, which strongly suggested that DNALI1 would have a function related to that of PACRG. Testing this hypothesis through the use of conditional *Dnali1* knockout mice revealed a co-dependence between the two proteins for structural integrity of spermatid elongation, the absence of which leads to abnormal sperm morphology, immotility and complete male infertility.

It is not surprising to see dramatically reduced sperm counts associated with impaired spermiogenesis, because DNALI1 forms a complex with MEIG1/PACRG. Disruption of either *Meig1* or *Pacrg* in the male germ cells resulted in a similar phenotype of impaired spermiogenesis and male infertility (26, 27, 34–38). The manchette is believed to play an important role in transporting cargo proteins for the formation of sperm flagella, and the transport function needs motor proteins (30, 40). Several motor proteins have been reported to be present in the manchette (41–44), and some have been shown to be essential for flagella formation (45, 46). It is highly possible that DNALI1 is the major driving force in transporting the MEIG1/PACRG complex together with the sperm flagellar proteins, including SPAG16L, along the manchette microtubules in order to assemble a functional sperm tail. It appears that DNALI1 is an upstream protein of MEIG1/PACRG complex, because DNALI1 is still in the manchette when PACRG is absent (Figure 10A). In the absence of DNALI1, MEIG1 and the cargo protein SPAG16L are no longer present in the manchette. As a light chain protein, it is unlikely that DNALI1 binds directly to the manchette microtubules, based on its localization in the transfected mammalian cells. However, DNALI1 has been reported to be a binding partner of dynein heavy chain 1 protein (DYNC1H1) (22) and our current study revealed evidence that the dynein heavy chain 1 protein is also present in the manchette (Supplemental Figure 7). Thus, it is more likely that DNALI1 associates with the manchette microtubules through dynein heavy chain 1. Another function of the manchette is to help shape the head of developing spermatids (47, 48). Like the *Meig1* and *Pacrg* mutant mice, the *Dnali1* cKO mice also developed abnormal sperm heads, which further supports the claim that DNALI1 is involved in the construction of a functional manchette.

**Figure 10.**
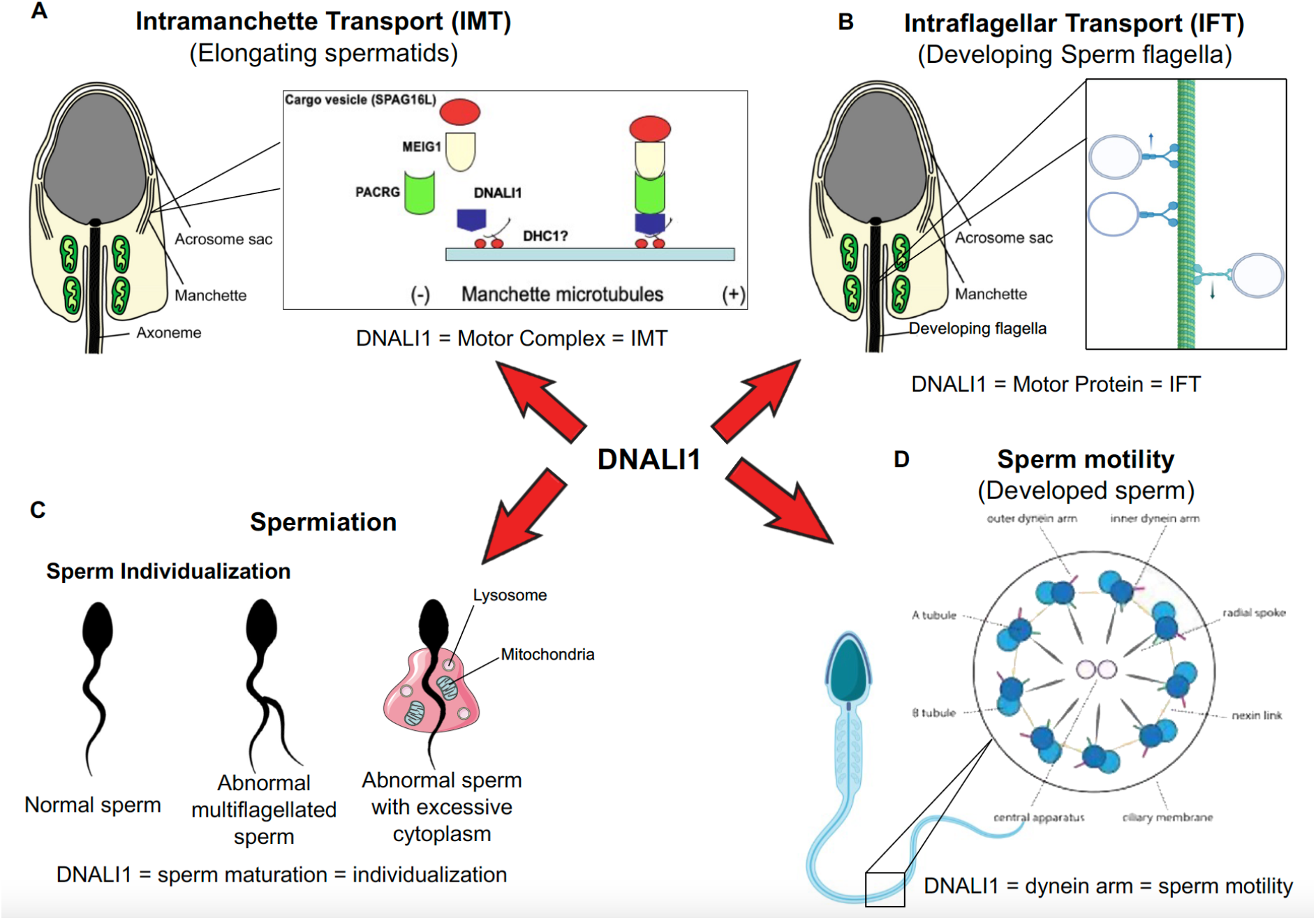
Working model of DNALI1 in sperm cell differentiation and function. A. DNALI1 forms a complex with MEIG1/PACRG, with DNALI1 being an upstream protein that recruits downstream PACRG and MEIG1 to the manchette. DNALI1 associates with the manchette microtubules through other molecular motor protein(s), including dynein heavy chain 1. MEIG1/PACRG/DNALI1/motor complex transports cargos, including SPAG16L along the manchette to build sperm flagella. B. DNALI1 might also function as a motor protein involved in transporting IFT particles. C. DNALI1 may facilitate in the appropriate maturation and individualization of sperm cells. D. DNALI1 is present in the dynein arm and functions in sperm motility.

Formation of the flagellum involves both the IMT and the IFT and the complex abnormal sperm phenotype observed in the *Dnali1* cKO appears to be due to the disruption of both pathways. The IMT mediates the transport of cargo proteins to the basal bodies, the template and the start point from which the sperm axoneme is formed (30). In the *Dnali1* cKO mice, although some sperm were formed, none had normal morphology. Besides abnormal heads, the flagella showed multiple defects, including short tails and vesicles and gaps along the sperm tail. The partial formation of the tails indicates that the IMT may not always be disrupted, possibly due to an incomplete excision by the Cre recombinase, allowing for limited DNALI1 protein synthesis.

The IFT is one of the more important mechanism involved in the formation of cilia/flagella (49). The IFT complex is composed of core IFT components, BBSomes, the motor proteins and cargo proteins (50) and the IFT transport process is bidirectional. Through antegrade transport, the cargo proteins are transported from the basal bodies to the tips of cilia/flagella and by the retrograde transport it is possible for the turn-over products to be transported back to the cell body for recycling. The morphological abnormalities observed in cKO sperm lead us to hypothesize that DNALI1 may also be involved in the intraflagellar transport (IFT) process. Defects in IFT result in failure of ciliogenesis, including formation of the sperm flagellum (51–58). Motor proteins have been shown to be essential for a functional IFT (59, 60). Therefore, given the ultrastructural abnormalities in the sperm flagella, loss of DNALI1 might also result in IFT dysfunction (Figure 10B), and be a major contributing factor to sperm immotility. Although we cannot exclude the possibility that other transport systems mediate the transport of cargo proteins in the manchette, data presented here show that the MEIG1/PACRG/DNALI1 complex does transport some cargo proteins that are essential for the formation of a normal sperm tail.

TEM study revealed a disorganized fibrous sheath, sperm tail structure in the *Dnali1* cKO mice. The fibrous sheath is a cytoskeletal structure of the principal piece in the flagellum, which consists of two longitudinal columns connected by semicircular ribs. The longitudinal columns are attached to outer dense fibers 3 and 8 in the anterior part of the principal piece and replace those fibers in the middle and posterior part of the principal piece and become associated with microtubule doublets 3 and 8 (61). It is assumed that such an elaborate substructural organization of the fibrous sheath into longitudinal columns and semicircular ribs must have a function, but testable hypotheses surrounding these characteristics are lacking. In *Dnali1* mutant mice, the two longitudinal columns were not associated with microtubule doublets 3 and 8, and the two semi circumferential ribs showed defective or asymmetrical alignment in the principal piece. It is speculated that the fibrous sheath serves as a scaffold for proteins participating in glycolysis, cAMP-dependent signaling transduction and mechanical support (62, 63). The cAMP-signaling pathway has been linked to the regulation of sperm maturation, motility, capacitation, hyperactivation, and the acrosome reaction (64). Therefore, defects in the development and function of the fibrous sheath may also contribute to infertility of *Dnali1* cKO mice. Dysfunction in IMT and IFT may contribute to these structural defects.

Another interesting phenotype observed was an occasional failure of sperm individualization, or disengagement (65–67), at the late spermatid step in the *Dnali1* cKO mice. This abnormality was not observed in the *Meig1* and *Pacrg* mutant mice (26, 27, 34–38). Individualization defects are the most common cause of human male infertility (68). Disengagement is one of the later steps of spermiogenesis, that occurs when the membrane-cytoskeleton individualized complex (IC) is assembled around the nucleus of each bundle of 64 haploid elongated spermatid nuclei. Each IC is composed of 64 F-actin-based investment cones, which move the flagella downward as a coordinated ensemble. Each spermatid is individualized of a single cone. As IC progresses, the cytoplasm is removed from the flagella, and the membrane surrounding each spermatid is reshaped to form the individualized spermatozoa (69, 70). It has been reported using genetic studies that at least 70 genes are involved in the process of disengagement and individualization. Several molecular pathways contribute to this process, including actin and microtubule dynamics, the ubiquitin-proteasome pathway components, apoptotic elimination of cytoplasmic contents, plasma membrane reorganization, and the formation of a disengagement complex (65, 67, 71–73). The microtubule cytoskeleton is important for individualization (73). Complexes of microtubules and dynein and kinesin motors form the core of cilia and flagella (74). Mutations in the components of the dynein-dynein complex, including the cytoplasmic dynein intermediate chain (CDIC), the two Drosophila dynein light chains DDLC1 and DLC90F, disrupt the synchronous movement of actin cones, but they also disrupt the nucleus shaping and positioning (75–77). These consistent phenotypes suggest that DNALI1 may have a similar effect on the synchronous movement of actin cones in sperm individualization and that DNALI1 has other functions in male germ cell development that differ from the MEIG1/PACRG complex (Figure 10C).

Another potential function of DNALI1 is its reported activity as an inner dynein arm (IDA) component of the cilium and flagellum axoneme (78). This would not be surprising in sperm, as it is present along the entire length of the sperm flagellar axoneme (22). Dynein proteins are present in the outer (ODAs) and inner dynein arms (IDAs) of the axonemal complex, and both arms are essential for the beating of cilia and flagella (78, 79). IDAs have seven major subspecies and four minor subspecies (60), but little is known about the functional differences between these subspecies (80–82). Therefore, if DNALI1 does serve this function in the sperm axoneme, its inactivation in male germ cells would likely disrupt the function of the dynein arms, and contribute to the formation of immotile sperm (Figure 10D).

In summary, DNALI1 potentially has multiple roles in sperm formation and function. We demonstrated a main function in IMT but additional roles of DNALI1 are possible in IFT and sperm individualization and disengagement, all of which being biological processes that occur during spermiogenesis. Failure of these processes causes impaired sperm formation and function and finally results in male infertility. While we show that an IMT function is clearly associated with the MEIG1/PACRG complex, the potential function of DNALI1 in IFT, sperm individualization and spermiation are not clear and will need further investigation.

## Acknowledgments

This research was supported by Wayne State University Start-up fund and Wayne State University Research Fund, MCI pilot award (to ZZ), MCI fellowship (to YTY), NIH R01DK76229 (JGG) and an NIH grant R01DK115563 (to DCW). Funding was also obtained from Agence Nationale pour la Recherche (grant FLAGELOME ANR-19-CE17-0014 to AT). We also thank Dr Scott C. Henderson and Frances K. White for their assistance with using the confocal microscopy in Microscopy Core Facility of Virginia Commonwealth University.

## Declaration of interest

There is no conflict of interest that could be perceived as prejudicing the impartiality of the research reported.

## Supplemental materials and legends

**Supplemental Figure 1.**
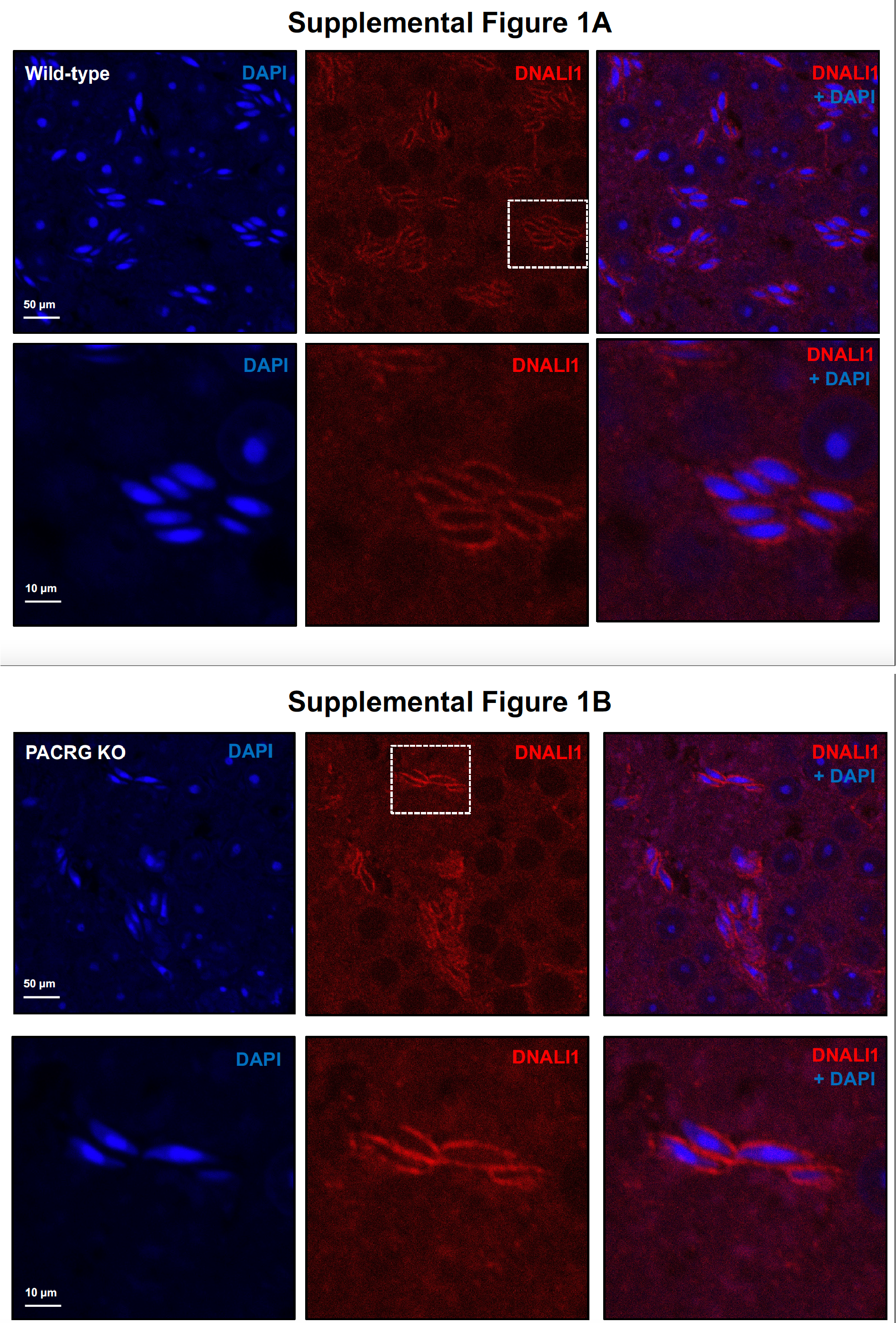
The localization of DNALI1 in the testis seminiferous tubule of a wild-type mouse (A) and a *Pacrg* mutant mouse (B). DNALI1 was present in the manchette of elongating spermatids, and the localization was not changed in the *Pacrg* mutant mice. The lower panels were the zoom-in areas from the upper panels. (Images taken with confocal laser scanning microscopy (Zeiss LSM 700), Virginia Commonwealth University).

**Supplemental Figure 2.**
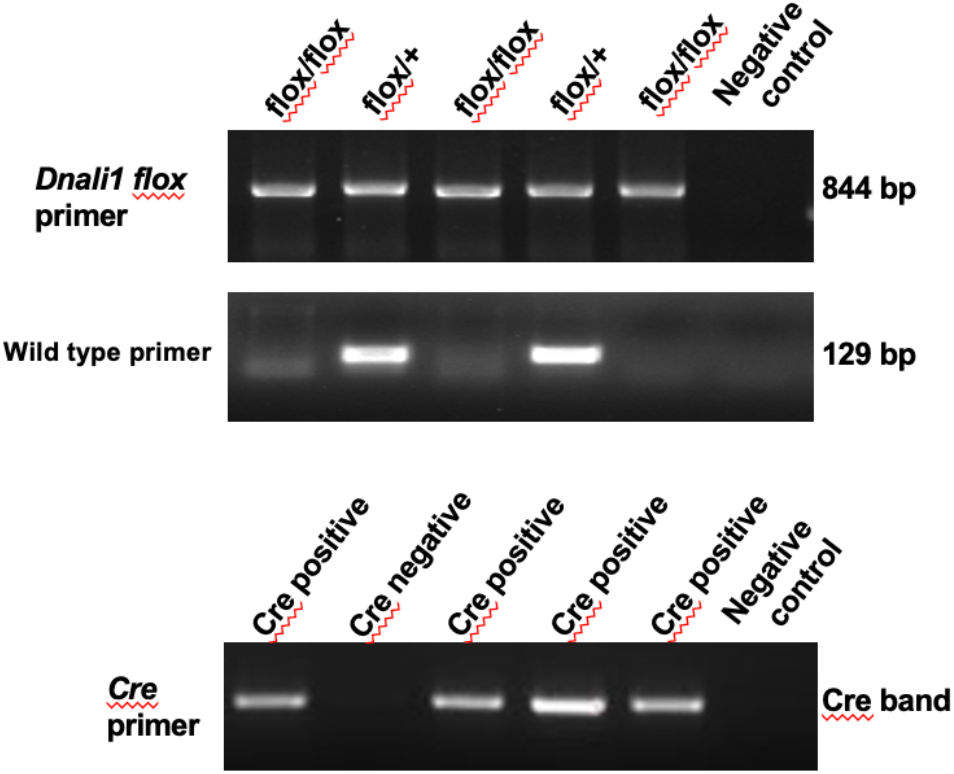
Representative PCR results showing mice with different genotypes. Upper panel: primer set to analyze floxed *Dnali1* allele (844 bp); middle panel: primer set to analyze the wild-type allele (129 bp); lower panel: primer set to detect Cre.

**Supplemental Figure 3.**
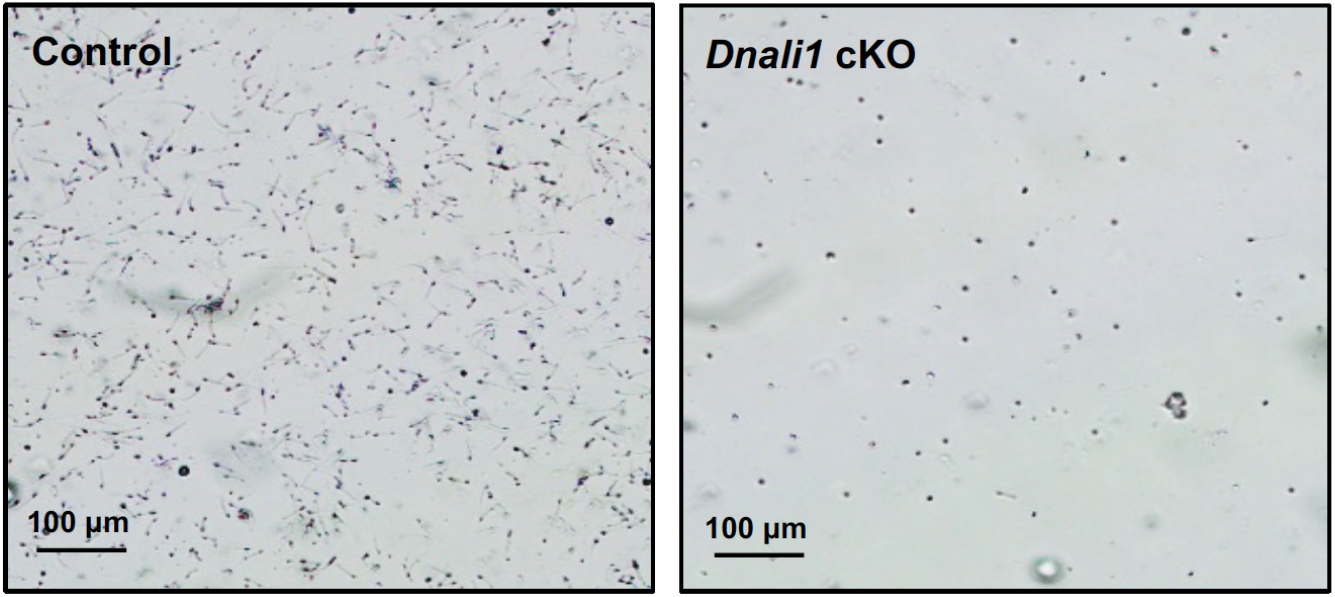
Morphological examination of epididymal sperm by light microscopy at low magnification. Notice that sperm density of the control mice is higher than those observed in the *Dnali1* cKO mice under the same dilution.

**Supplemental Figure 4.**
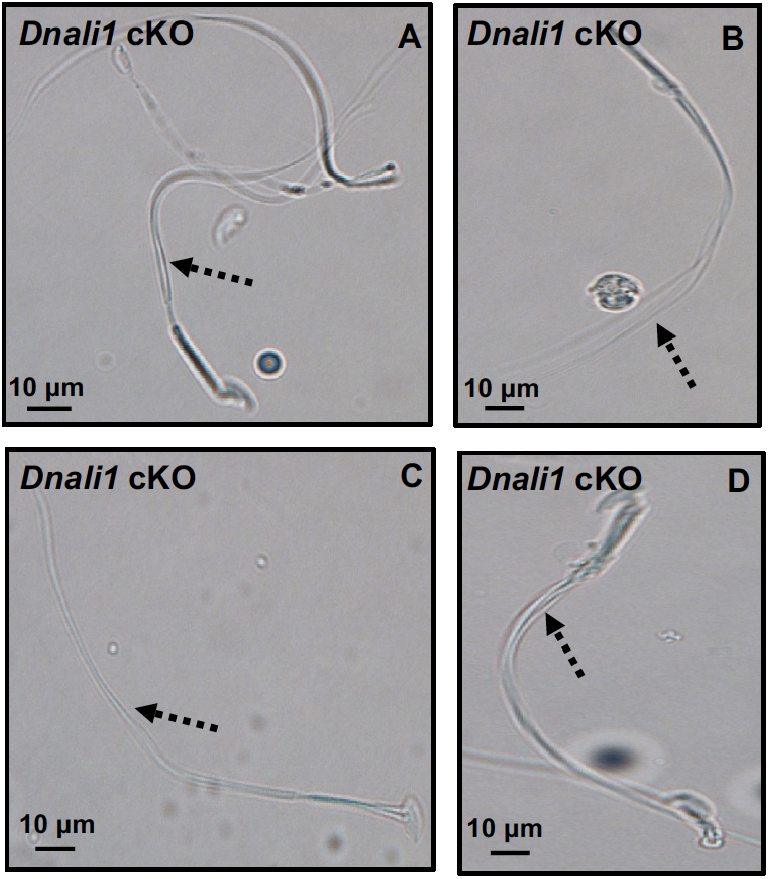
Morphological examination of epididymal sperm by light microscopy. Multiple tails are present in the sperm (dashed arrows in A-D).

**Supplemental Figure 5.**
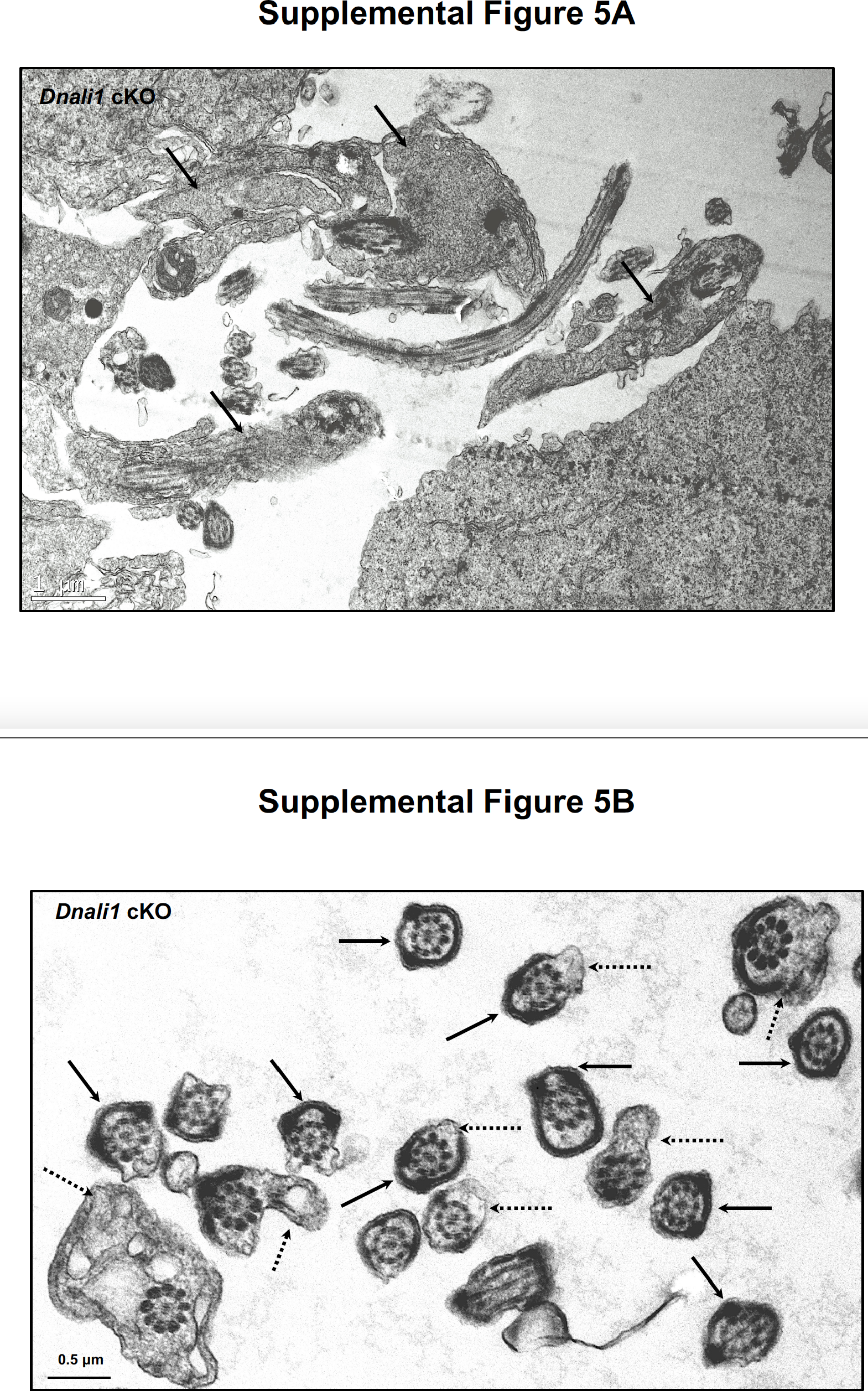
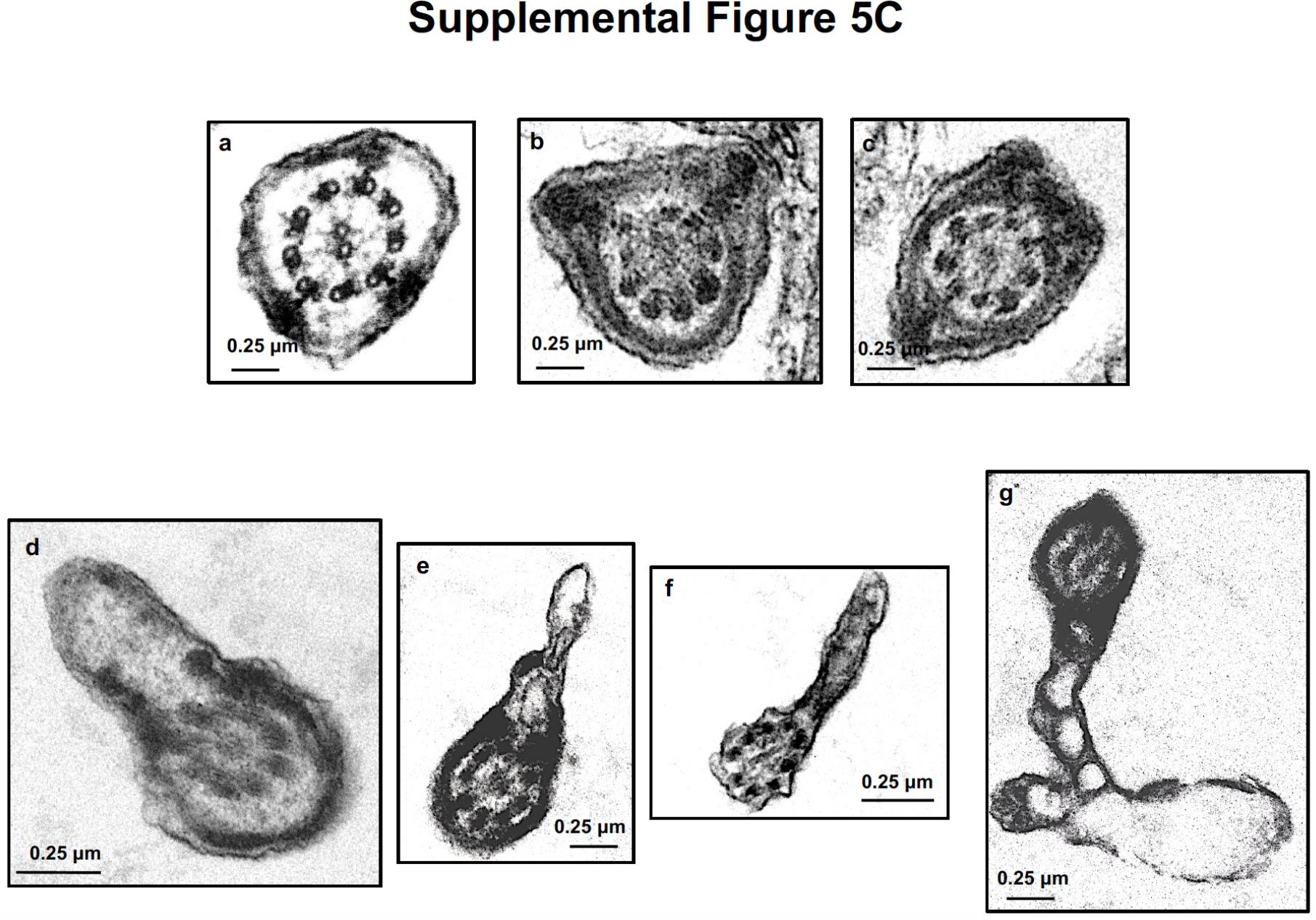
Additional testicular sperm TEM images of the *Dnali1* cKO mice. A. Low magnification image showing retained cytoplasmic components in the lumen area (black arrows). B. Low magnification images showing abnormal fibrous sheath (black arrows) and cell membranes (dashed arrows); C. High magnification images showing abnormal fibrous sheath (a-c) and cell membranes (d-g).

**Supplemental Figure 6.**
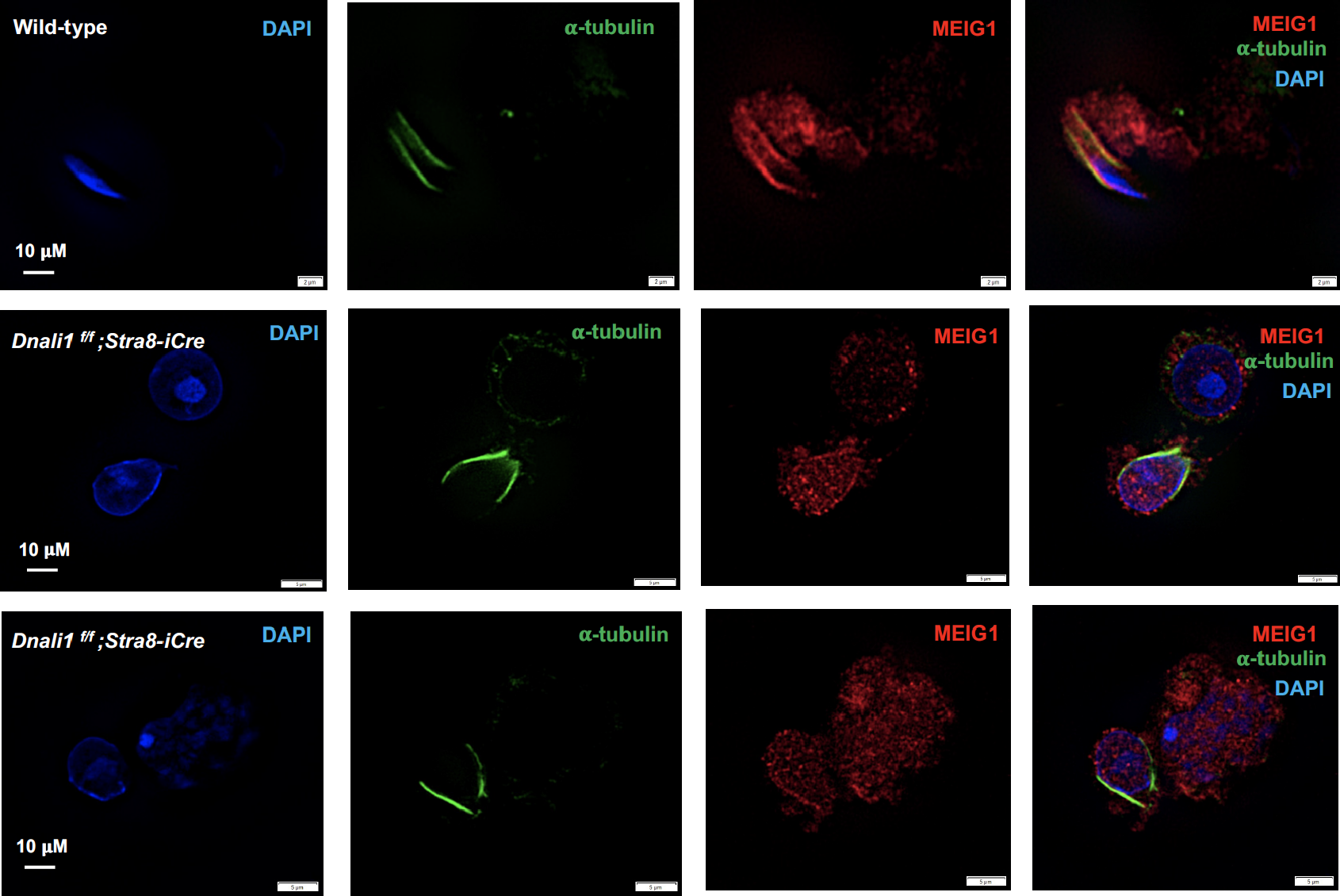
Examination of MEIG1 localization in elongating spermatids in the control and *Dnali1* cKO mice. Notice that MEIG1 is present in the manchette in the control mouse, but not in the *Dnali1* cKO mouse. (Images taken with Olympus IX-81 microscope equipped with a spinning-disc confocal unit (Dr. James G. Granneman’s laboratory, Wayne State University).

**Supplemental Figure 7.**
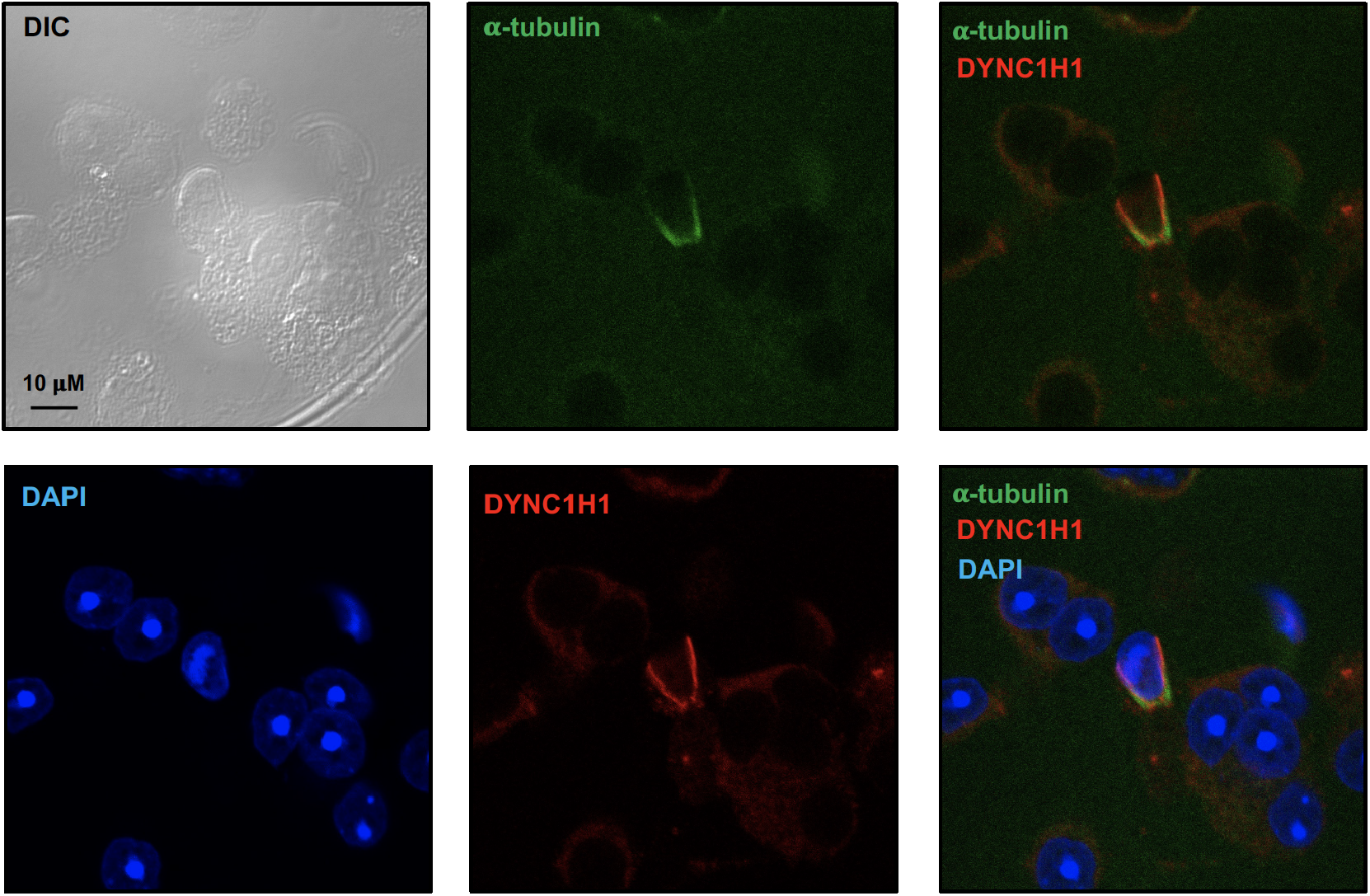
Examination of dynein heavy chain 1 protein (DYNC1H1) in male germ cells by immunofluorescence staining. Notice that DYNC1H1 was co-localized with α-tubulin in the elongating spermatid. (Images taken with confocal laser scanning microscopy (Zeiss LSM 700), Virginia Commonwealth University).

**Supplemental Table 1.** **List of putative PACRG binding proteins selected under stringent conditions.** The full-length PACRG coding sequence was cloned into pGBKT7, which was used to screen a Mate & PlateTM Library-Universal Mouse (Normalized) (Clontech, Cat#: 630482) according to the manufacturer’s instructions. The yeasts were grown on plates lacking four amino acids (Ade-Leu-His-Trp). DNALI1 was found to be one of the putative PACRG binding proteins.

**List of putative PACRG binding proteins selected under stringent conditions**
| Name | NCBI number | Frequency |
| --- | --- | --- |
| Meig1 | NM_008579 | 119 |
| **Dnali1** | **NM_175223** | **6** |
| Pramel42 | NM_001243938 | 3 |
| Musculus protein phosphatase 1A | BC008595 | 2 |
| Acad11 | NM_175324 | 2 |
| Ppm1a | NM_008910 | 1 |
| L2hgdh | NM_145443 | 1 |
| Tmem225 | NM_029379 | 1 |
| Tinag | NM_012033 | 1 |
| Spag6l | NM_015773 | 1 |
| Emp2 | NM_007929 | 1 |

## Supplemental movies

A. Representative movie from a control mouse. Note that most sperm are motile and display vigorous flagellar activity and progressive long-track forward movement.

B. Representative movies from a *Dnali1* cKO mutant mouse. Notice that there are fewer sperm compared with the control mice in the same dilution, and almost all sperm are immotile.

## Notes

### Competing Interest Statement

The authors have declared no competing interest.

